# Non-Necroptotic Roles of MLKL in Diet-Induced Obesity, Liver Pathology, and Insulin Sensitivity: Insights from a High Fat, High Fructose, High Cholesterol Diet Mouse Model

**DOI:** 10.1101/2024.01.10.575102

**Authors:** Phoebe Ohene-Marfo, Hoang Van M Nguyen, Sabira Mohammed, Nidheesh Thadathil, Albert Tran, Evan H Nicklas, Dawei Wang, Ramasamy Selvarani, Jacob Farriester, Rohan Varshney, Michael Kinter, Arlan Richardson, Michael Rudolph, Sathyaseelan S. Deepa

## Abstract

Chronic inflammation is a key player in metabolic dysfunction-associated fatty liver disease (MAFLD) progression. Necroptosis, an inflammatory cell death pathway, is elevated in MAFLD patients and mouse models, yet its role is unclear due to diverse mouse models and inhibition strategies. In our study, we inhibited necroptosis by targeting mixed lineage kinase domain like pseudokinase (MLKL), the terminal effector of necroptosis, in a high-fat, high-fructose, high-cholesterol (HFHFrHC) mouse model of diet-induced MAFLD mouse model. Despite HFHFrHC diet upregulating MLKL (2.5-fold), WT mice livers showed no increase in necroptosis markers or associated proinflammatory cytokines. Surprisingly, *Mlkl^−/−^*mice experienced exacerbated liver inflammation without protection from diet-induced liver damage, steatosis, or fibrosis. In contrast, *Mlkl^+/−^* mice showed significant reduction in these parameters that was associated with elevated Pparα and Pparγ levels. Both *Mlkl^−/−^*and *Mlkl^+/−^* mice on HFHFrHC diet resisted diet-induced obesity, attributed to increased beiging, enhanced oxygen consumption and energy expenditure due to adipose tissue, and exhibited improved insulin sensitivity. These findings highlight the tissue specific effects of MLKL on the liver and adipose tissue, and suggest a dose-dependent effect of MLKL on liver pathology.

## INTRODUCTION

Metabolic associated fatty liver disease (MAFLD) is a prevalent global liver condition linked to obesity, type 2 diabetes, and insulin resistance (1). MAFLD covers a spectrum of liver diseases ranging from fat deposition in the liver (steatosis) to metabolic dysfunction-associated steatohepatitis (MASH), which is characterized by fat accumulation (steatosis) in *>*5% of hepatocytes, and inflammation with hepatocyte injury (ballooning) with varying degrees of fibrosis (2). The obesity epidemic has led to a rapid increase in MAFLD prevalence, affecting nearly 38% of the population (3). Around 30% of those with steatosis progress to MASH, a significant cause of liver-related morbidity and mortality. MASH is a leading factor in liver transplantation cases and is a major risk factor for hepatocellular carcinoma (HCC) and cardiovascular diseases (4, 5). Despite ongoing clinical trials, there is currently no approved pharmacological approach against NASH, emphasizing the need for a deeper understanding of its pathogenic mechanisms.

One of the factors that differentiates MASH from simple fatty liver is the occurrence of massive hepatocyte cell death in MASH and subsequent increase in liver inflammation and fibrosis (6). Hepatocyte cell death is a critical event in the development and progression of MAFLD because it triggers inflammation leading to fibrosis. Apoptosis is a major form of cell death in liver diseases, however, in recent years, necroptosis has emerged as an important form of programmed cell death in the liver (7). Apoptotic cell death is characterized by lower levels of inflammation, whereas necroptosis induces significant inflammation due to the release of damage-associated molecular patterns (DAMPs). Necroptosis is initiated when necroptotic stimuli (e.g. oxidative stress, TNFα, mTOR activation, lipotoxicity etc.) sequentially activate receptor interacting serine/threonine kinase 1 (RIPK1), RIPK3, and mixed lineage kinase domain like pseudokinase (MLKL) through phosphorylation. Phosphorylated MLKL undergoes oligomerization leading to membrane permeabilization leading to the release of DAMPs, which bind to cell surface receptors on innate immune cells leading to increased transcription of proinflammatory cytokines causing inflammation (8, 9). Studies have shown that markers of necroptosis (RIPK3, MLKL, and phospho-MLKL) are increased in the livers of MAFLD and MASH patients, and in mouse models of MASH (10–15). Consequently, inhibiting necroptosis, either through pharmacological targeting of RIPK1 or genetic/pharmacological targeting of RIPK3, has been demonstrated to decrease liver inflammation and fibrosis in mouse models of MAFLD (12, 16, 17). We have previously demonstrated that the inhibition of necroptosis using necrostatin-1s, a pharmacological inhibitor of RIPK1, reduced liver inflammation and fibrosis in mouse models of spontaneous MASH, i.e. old wild type mice and *Sod1^−/−^*mice (18, 19). However, conflicting reports exist in the literature regarding the impact of *Ripk3* or *Mlkl* deletion on liver inflammation and/or fibrosis, with outcomes varying depending on the dietary model of MAFLD.

In a methionine choline-deficient (MCD) or choline-deficient high-fat diet (CD-HFD, 35% fat) non-obese model, RIPK3 deletion reduces steatosis, inflammation, and fibrosis (12, 14). Conversely, in a HFD-induced obesity-driven MAFLD, RIPK3 deletion exacerbates outcomes, including increased body weight, insulin resistance, glucose intolerance, and heightened liver steatosis, inflammation, and fibrosis (13, 20). Thus, RIPK3’s role in liver inflammation and fibrosis is model-dependent and influenced by diet composition. Similarly, MLKL loss also yielded varied outcomes in MAFLD mouse models: *Mlkl* deletion reduced liver inflammation in HFD or western diet-induced MAFLD mouse models (15, 21), however, Xu et al (2018) reported no impact of *Mlkl* deletion on liver inflammation in an HFD-induced model of MAFLD (22). Notably, the effect of targeting *Mlkl* on liver fibrosis in diet-induced MAFLD mouse models is unclear. Recently, we have shown that in a CD-HFD (60% fat) non-obese model, either *Ripk3* or *Mlkl* deletion reduced liver inflammation and HCC, whereas liver fibrosis was not reduced (23).

In the present study, we investigated the impact of the absence of *Mlkl* (*Mlkl^−/−^* mice) or partial loss of *Mlkl* (*Mlkl^+/−^*mice) on MAFLD using a high-fat, high-fructose, high-cholesterol (HFHFrHC) diet that promotes obesity, fibrosis, and associated metabolic disorders in mice (24). Our data show that absence of MLKL did not protect against liver injury or fibrosis induced by the HFHFrHC diet. However, a partial loss of *Mlkl* protected from HFHFrHC diet induced liver injury and fibrosis. Interestingly, both *Mlkl^−/−^* and *Mlkl^+/−^*mice displayed resistance to HFHFrHC diet-induced obesity and improved insulin sensitivity compared to control wild type mice (WT, *Mlkl^+/+^*) fed the same diet.

## RESULTS

### *Mlkl^−/−^* and *Mlkl^+/−^* mice are resistant to diet induced obesity

The experimental design is shown in **Fig. 1A** where WT, *Mlkl^−/−^* and *Mlkl^+/−^* male mice were fed a HFHFrHC diet for 6 months, starting at 2 months of age. Feeding a HFHFrHC resulted in a significant increase (15.4%) in total body weight. Notably, this weight gain was not observed in *Mlkl^−/−^*or *Mlkl^+/−^* mice (**Figs. 1B, S1A**). Compared to Control Diet (CD, open bars) fed WT mice, HFHFrHC diet (grey bars) fed WT mice had a 2.6-fold increase in percentage fat mass. However, either absence or partial loss of *Mlkl* (*Mlkl^−/−^* or *Mlkl^+/−^*) led to significant attenuation of fat deposition (**Fig. 1C**). The lean mass to fat mass ratio was significantly lower in WT mice fed a HFHFrHC diet, whereas this effect was attenuated in *Mlkl^−/−^* or *Mlkl^+/−^* mice (**Fig. 1D**). No significant differences in food or water intake were observed between WT or *Mlkl^−/−^* mice provided either CD or HFHFrHC (**Figs. 1E, S1B-S1D**). CD-fed *Mlkl^−/−^* mice exhibited a significant reduction in ambulatory activity during dark phase compared to both CD-fed or HFHFrHC diet-fed WT groups (**Figs. 1F, S1E**). Together, based on increased body weight and adipose accumulation, the diet-induced obesity in HFHFrHC diet fed WT mice is attenuated in *Mlkl^−/−^* and *Mlkl^+/−^* mice and is not accounted for by changes in energy intake or ambulatory activity.

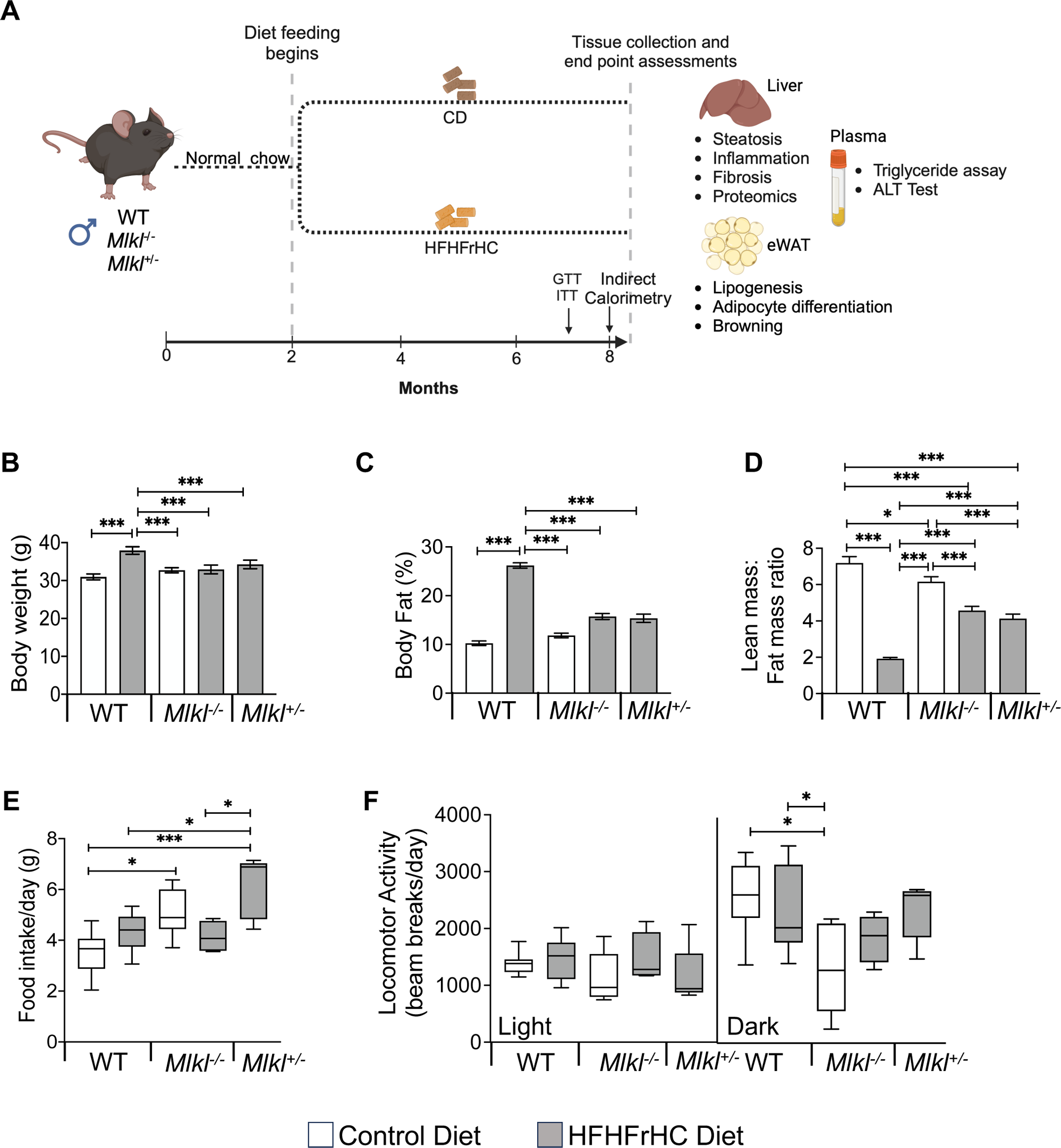

### *Mlkl^+/−^* mice, not *Mlkl^−/−^* mice, are protected from diet-induced hepatic injury and steatosis

HFHFrHC diet feeding resulted in a significant increase in the percentage liver weights of WT, *Mlkl^−/−^* and *Mlkl^+/−^*mice (∼1.3-fold) (**Fig. 2A**). As anticipated, consumption of HFHFrHC diet resulted in a significant increase in the MLKL protein (2.5-fold) in WT mice. However, unexpectedly, a 50% reduction in RIPK3 protein and no change in phospho-MLKL and MLKL oligomers (markers of necroptosis) were observed (**Figs. S2A, S2B**). Feeding a HFHFrHC diet increased the levels of triglycerides in the liver (2-2.5-fold) (**Fig 2B**). H&E staining revealed accumulation of enlarged lipid droplets in the liver of WT mice fed a HFHFrHC diet compared to WT mice fed CD, indicating increased steatosis (**Figs. 2C, 2D**). *Mlkl^−/−^* mice fed the HFHFrHC diet group had smaller-sized lipid droplets in the liver, however, the number of lipid droplets were ∼2-fold greater than those found in either WT or *Mlkl^+/−^* HFHFrHC groups (**Figs. 2E**). In contrast, *Mlkl^+/−^* mice on the HFHFrHC diet had fewer and smaller-sized lipid droplets relative to the WT HFHFrHC group (**Figs. 2B-E**). Plasma triglyceride levels increased 1.5-fold, as did plasma ALT levels (2-fold) in the WT HFHFrHC group (**Figs. 2F, 2G**). Absence of MLKL (*Mlkl^−/−^* mice) did not reduce plasma TAG or ALT levels, however, partial loss of MLKL (*Mlkl^+/−^* mice) had significantly lower plasma triglycerides and ALT (**Figs. 2F, 2G**). In line with this observation, the transcript levels of Pparα and Pparγ, two key regulators of lipid metabolism (25, 26), were significantly elevated in the livers of *Mlkl^+/−^* mice compared to the other experimental groups (**Fig. 2H**).

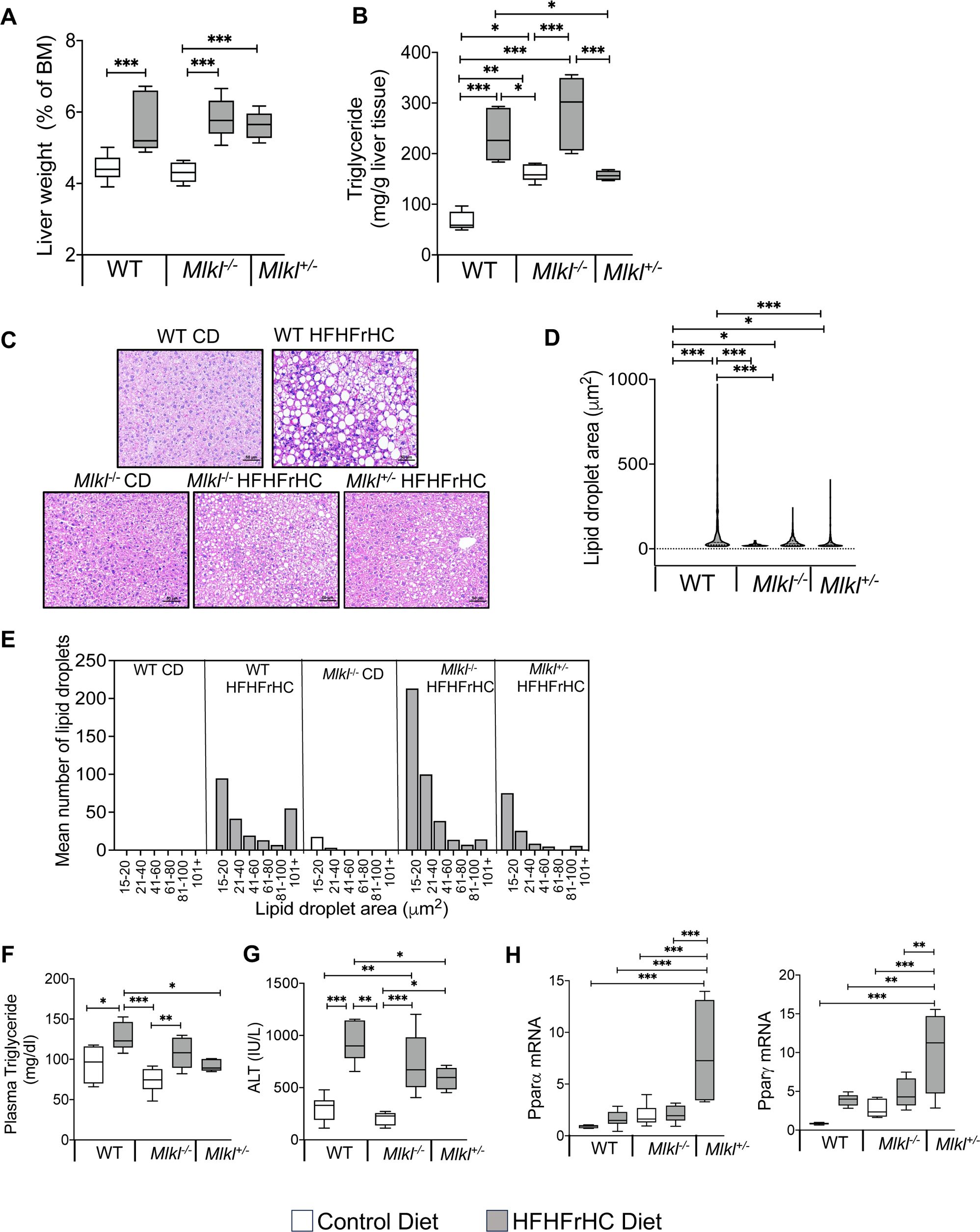

Next, we conducted targeted proteomic analysis to assess the effect of absence or reduction of *Mlkl* on proteins involved in mitochondrial fatty acid oxidation enzymes (**Fig. 3A**). The WT HFHFrHC group had significant increases in the protein levels for key fatty acid oxidation pathway enzymes, including acetyl-CoA carboxylase alpha and beta (ACCA1a/b, facilitating fatty acid oxidation), acyl-CoA synthetase long-chain family member 1 (ACSL1, involved in activating long-chain fatty acids for beta-oxidation), medium-chain acyl-CoA dehydrogenase (ACADM, participating in fatty acid beta-oxidation), enoyl-CoA hydratase and 3-hydroxyacyl CoA dehydrogenase (EHHADH, contributing to fatty acid beta-oxidation), 3-hydroxybutyrate dehydrogenase 1 (BDH1, involved in ketone body metabolism), and 3-hydroxyacyl-CoA dehydrogenase (HADH, engaged in fatty acid beta-oxidation). While ACC1a/b, BDH1, and EHHADH were also significantly increased in the *Mlkl^−/−^* group during HFHFrHC feeding, abundance of BDH1 and EHHADH was decreased significantly in the *Mlkl^+/−^*HFHFrHC group, relative to the WT HFHFrHC group (except for ACCA1a/b). Cumulatively, the absence or partial loss of MLKL has distinct effects on proteins regulating fatty acid metabolism compared to mice with intact MLKL.

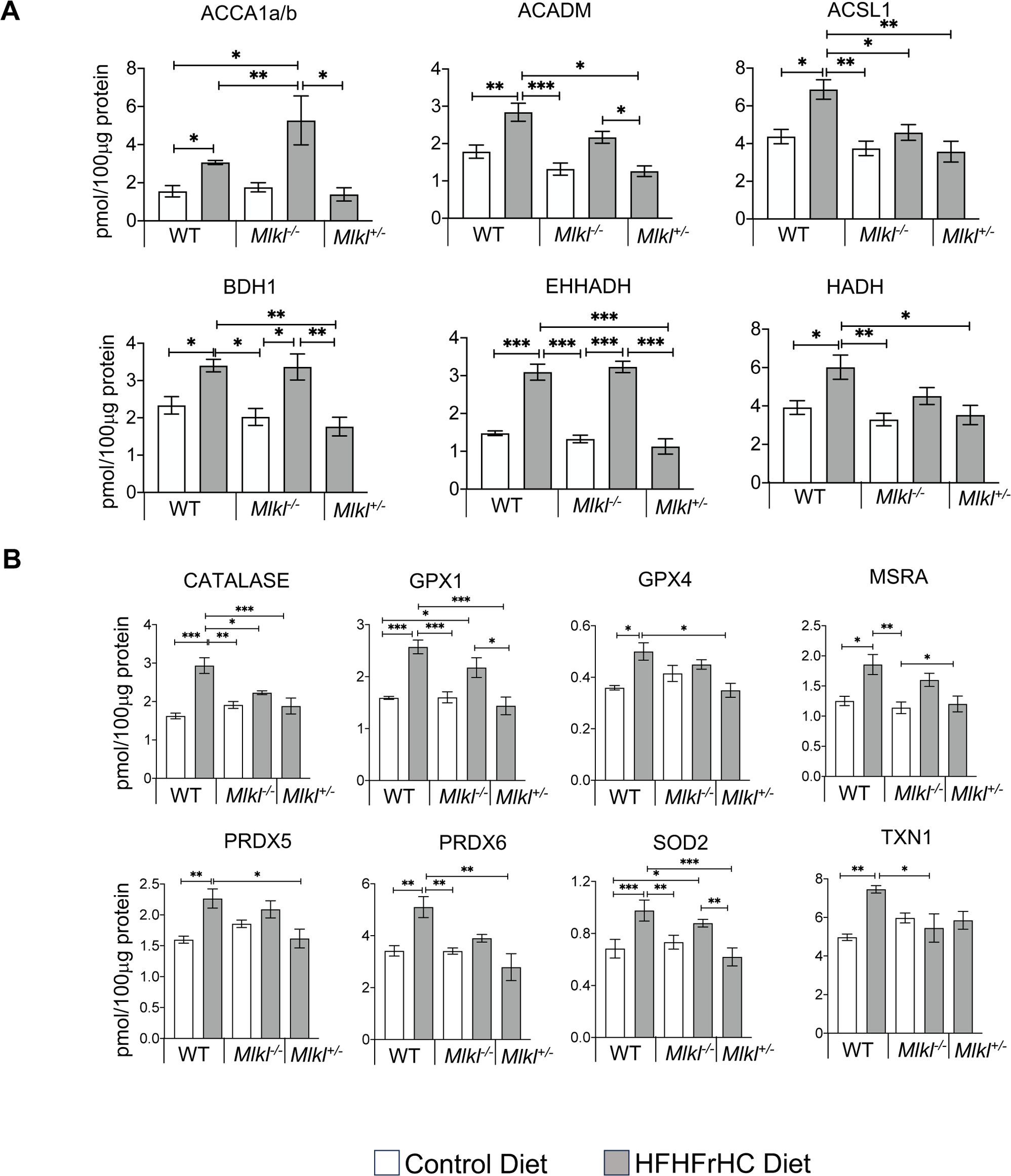

Fatty acid oxidation is known to elevate mitochondrial reactive oxygen species (ROS) levels, inducing oxidative stress (27), which is a key contributor to MAFLD development (28). Therefore, we evaluated antioxidant enzyme levels, as tissues upregulate these enzymes to protect against ROS and is a useful predictor of oxidative stress (29, 30). Proteomic analysis revealed significant increases in key antioxidant enzymes in response to HFHFrHC diet feeding in WT mice, including Mn-superoxide dismutase (SOD2, 1.6-fold), glutathione peroxidase (GPX1, 2.3-fold), GPX4 (1.6-fold), methionine sulfoxide reductase A (MSRA, 1.5-fold), peroxiredoxin 5 (PRDX5, 1.5-fold), PRDX6 (1.4-fold), thioredoxin 1 (TXN1, 1.6-fold), and catalase (2-fold) (**Fig. 3B**). In contrast, only SOD2 and GPX1 were significantly upregulated in the *Mlkl^−/−^*HFHFrHC group. On the other hand, partial loss of MLKL during HFHFrHC feeding restored levels of the mitochondrial antioxidant enzymes to those observed in the WT HFHFrHC group. Thus, the diminished fatty acid metabolism in HFHFrHC-fed *Mlkl^+/−^* mice was associated with decreased markers of oxidative stress. Collectively, these data indicate that under HFHFrHC diet conditions, absence or partial reduction of *Mlkl* has distinct effects on lipid droplet morphology and fat metabolism in the liver. Accordingly, only partial loss of MLKL conferred protection against HFHFrHC diet-induced liver damage and triglyceride accumulation.

### Absence of *Mlkl* exacerbated liver inflammation and had no impact on HFHFrHC diet-induced liver fibrosis

We next investigated the impact of absence or reduction of *Mlkl* on proinflammatory cytokines and chemokines in the liver because of the association of necroptosis with increased liver inflammation (19, 31) and the documented reduction of proinflammatory cytokines in mice with *Mlkl* deficiency on a western diet (21). The RNA expression of 84 proinflammatory cytokines and chemokines in the livers of WT, *Mlkl^−/−^*, and *Mlkl^+/−^* mice subjected to either a CD or HFHFrHC diet were quantified. The detailed data from the mouse cytokines and chemokines array are provided in **Table S2**. Feeding a HFHFrHC diet significantly upregulated 8 proinflammatory cytokines/chemokines in WT mice, and the top 5 genes elevated were Ccl22 (42-fold), IL-1rn (16-fold), IL-12b (15-fold), Ifna2 (12-fold), and IL-11 (5-fold) (**Table S2**). Interestingly, *Mlkl^−/−^*mice on HFHFrHC diet significantly elevated 44 proinflammatory cytokine/chemokines in the liver when compared to WT mice on CD, and the top 5 genes upregulated were Spp1 (53-fold), Ccl22 (46-fold), Lif (35-fold), Ccl11 (33-fold), and IL-1rn (21-fold). Importantly, necroptosis-associated proinflammatory cytokines TNFα (11-fold), IL6 (5.7-fold), IL-1β (4.6-fold), or CCL2 (15-fold) were also increased in the livers of *Mlkl^−/−^* fed a HFHFrHC diet (**Fig. 4A**) (**Table S2**). *Mlkl^+/−^* mice fed the HFHFrHC diet significantly upregulated 17 proinflammatory cytokine/chemokine genes and the top 5 genes were Ccl22 (96-fold), Xcl1 (36-fold), IL-12b (20-fold), Ifna2 (15-fold), and Csf3 (2-fold). Overall, unexpectedly, absence of *Mlkl* increased proinflammatory cytokines and chemokines in the livers of HFHFrHC-fed mice.

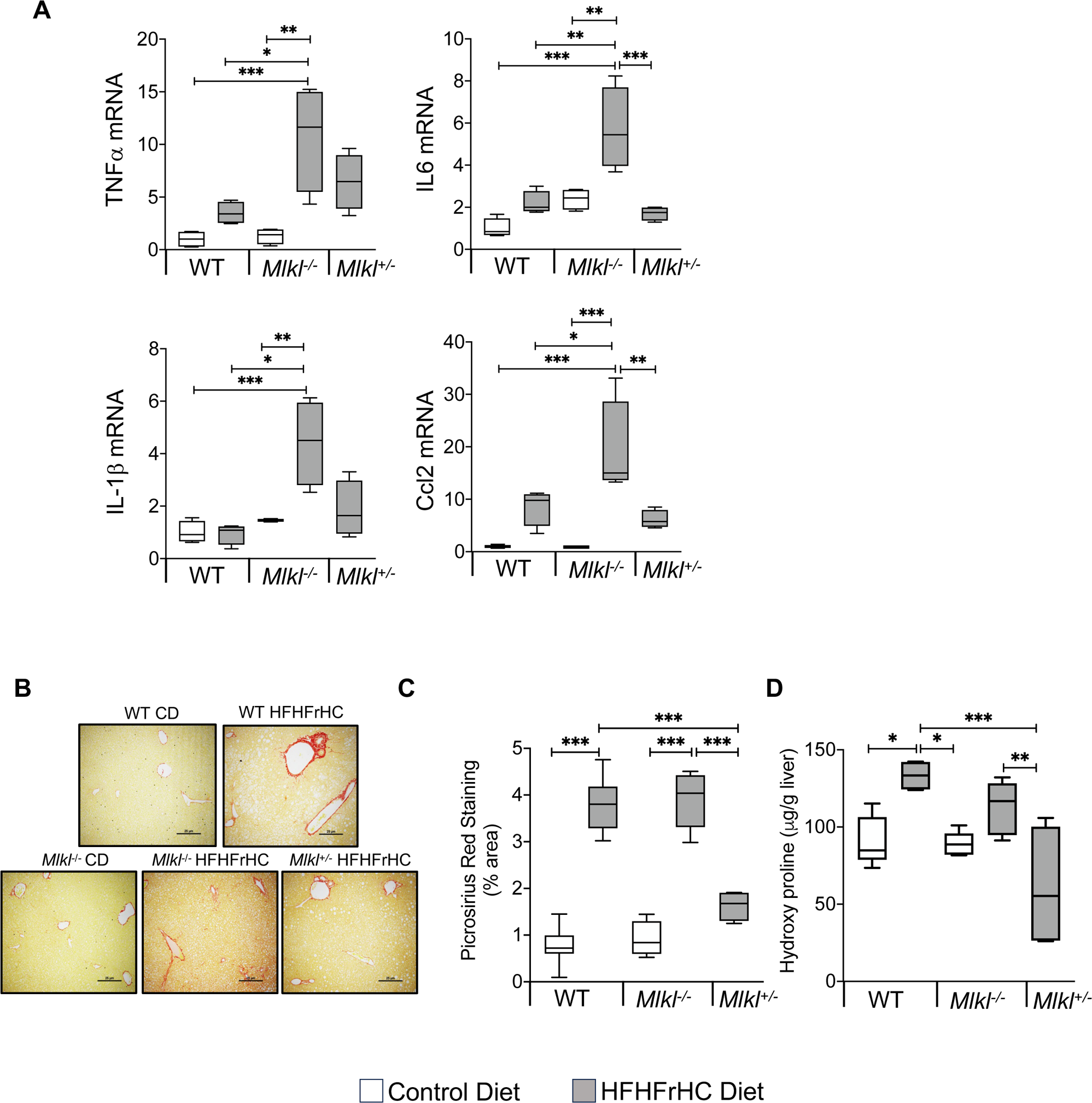

Hepatocyte injury, inflammation, and the innate immune system promote liver fibrosis through the activation of hepatic stellate cells and the secretion and deposition of extracellular matrix (32). Assessment of liver fibrosis revealed a significant increase in collagen fibers in the livers of WT mice fed the HFHFrHC diet as measured by PSR staining (3 to 3.5-fold) or hydroxyproline content (1.5 to 2-fold), compared to those on the CD (**Figs. 4B-4D**). While the absence of *Mlkl* did not attenuate liver fibrosis induced by the HFHFrHC diet, *Mlkl^+/−^* mice exhibited a significant reduction in fibrosis. Collectively, these data suggest that while the absence of *Mlkl* did not protect against liver fibrosis induced by HFHFrHC diet, partial loss of *Mlkl* is protective.

To further confirm the inability of *Mlkl^−/−^* mice to protect against liver fibrosis, we used a methionine choline-deficient (MCD), which is a widely used diet that induces fibrosis without obesity (33, 34). Our data in **Figs. S3A-C** show that feeding a MCD diet resulted in comparable levels of liver fibrosis markers [PSR staining (6-6.5-fold), hydroxyproline levels (1.8-fold), and fibrosis markers (Col1α1, 4-5-fold and Col3α1, 2-2.5-fold)] in WT and *Mlkl^−/−^* mice compared to mice fed a methionine choline-sufficient diet (MSD), control diet (CD). Transcript levels of TNFα, IL6, and IL1β were similar in MCD diet fed WT and *Mlkl^−/−^* mice, whereas Ccl2 levels were further exacerbated by the absence of *Mlkl* (**Fig. S3D**). Thus, absence of *Mlkl* is not protective against diet-induced liver fibrosis.

### *Mlkl^−/−^* and *Mlkl^+/−^* mice exhibited reduced adiposity in response to HFHFrHC diet

Contrary to the differential effects on the liver, both *Mlkl^−/−^* and *Mlkl^+/−^* mice on HFHFrHC diet showed similar resistance in diet-induced adipose accumulation (**Fig. 1C**). Consistent with their body composition data, feeding a HFHFrHC diet resulted in a significant increase (2.5-fold) in eWAT deposition, normalized to body weight, of WT mice but not for either *Mlkl^−/−^* or *Mlkl^+/−^* mice (**Fig. 5A**). Histological examination of eWAT showed that WT mice on a HFHFrHC diet had different adipose morphologies, and adipocyte cellularity analysis reveal that HFHFrHC *Mlkl^−/−^*and *Mlkl^+/−^* had significantly smaller adipocyte size distributions (**Figs. 5B, 5C**). Moreover, the average adipocyte area shifted towards smaller adipocytes in *Mlkl^−/−^* and *Mlkl^+/−^* mice fed a HFHFrHC diet compared to WT mice on the same diet (**Fig. 5D**). Next, we examined whether the decreased adipocyte size in *Mlkl^−/−^* or *Mlkl^+/−^* mice fed the HFHFrHC diet was due to impairments in either adipocyte differentiation or *de novo* lipogenesis. Absence of MLKL did not appear to impair the expression of key adipocyte differentiation regulators CCAAT enhancer binding protein alpha (Cebpα) or peroxisome proliferator-activated receptor gamma (Pparγ) in the eWAT of *Mlkl*^−/−^ and *Mlkl^−/−^* mice on a CD. In response to the HFHFrHC diet, Cebpα and Pparγ were suppressed in the WT group but significantly increased in both the *Mlkl^−/−^*and *Mlkl^+/−^* groups relative to the WT HFHFrHC mice (**Fig. 5E**). Analysis of *de novo* lipogenesis markers [fatty acid synthase (Fasn) and stearoyl-CoA desaturase (Scd)] revealed that absence of MLKL (*Mlkl^−/−^* mice) significantly elevated Scd levels but not FASN on a CD, whereas, feeding a HFHFrHC diet significantly reduced the levels of Fasn but not SCD in WT and *Mlkl^−/−^* mice. In contrast, both Fasn and Scd were significantly upregulated in *Mlkl^+/−^* mice (**Fig. 5F**). Analysis of transcript levels of proinflammatory cytokines/chemokines showed that feeding a HFHFrHC diet increased the level of Ccl2 (2.5-fold) but not TNFα, IL6, or IL1β, in the eWAT of WT mice, and Ccl2 levels were significantly downregulated in *Mlkl^−/−^* and *Mlkl^+/−^* mice fed the HFHFrHC diet (**Fig. S4A-4D**). These results suggest that the improved adipocyte cellularity sizes of the *Mlkl^−/−^* or *Mlkl^+/−^*mice during HFHFrHC feeding is due to enhanced expression of adipogenic regulators Cebpα and Pparγ.

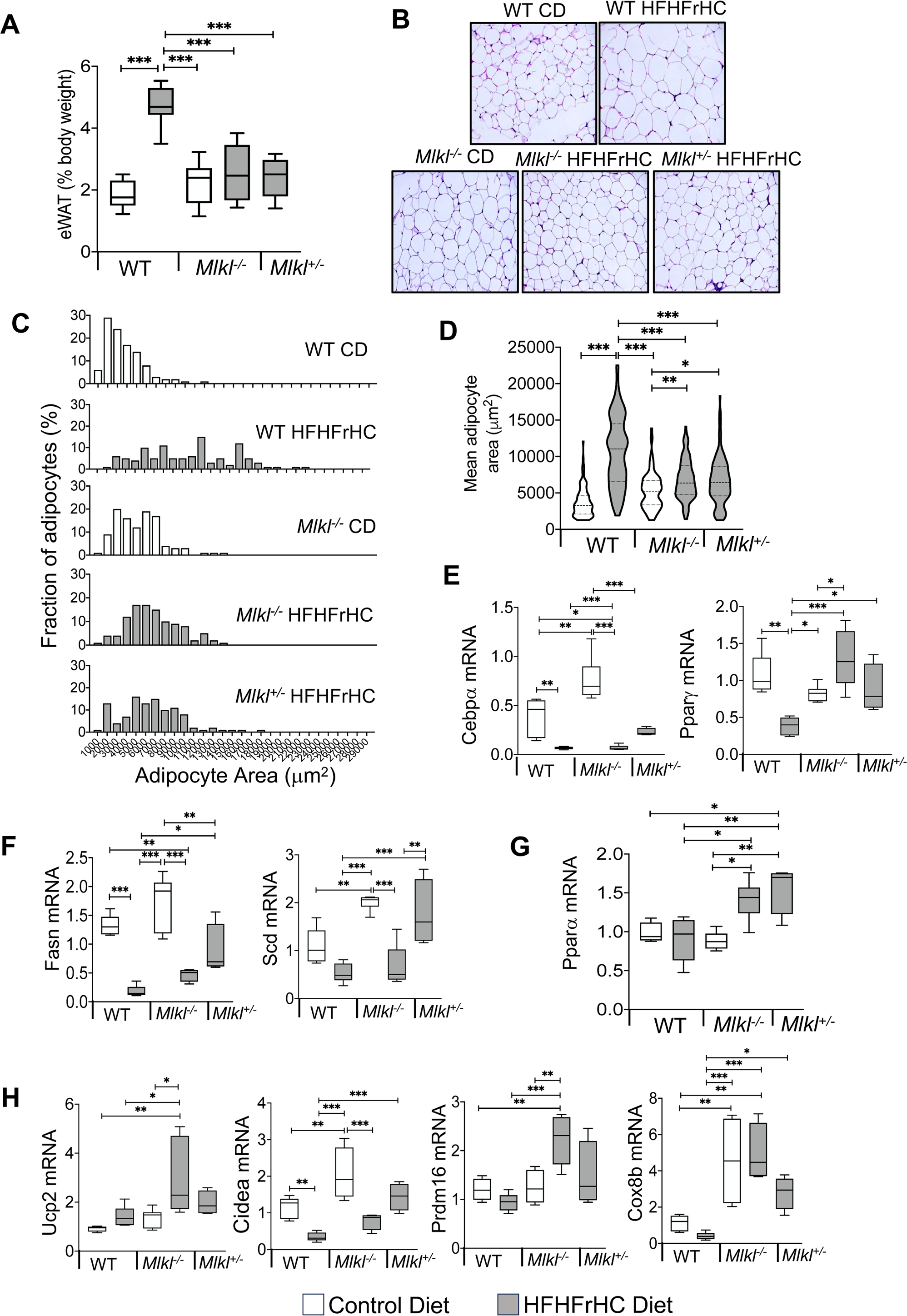

Next we tested whether levels of Pparα, a key regulator of lipid metabolism and energy expenditure in adipose tissue is altered in eWAT (35). The HFHFrHC diet did not alter Pparα levels in the eWAT of WT mice, whereas Pparα was significantly upregulated in *Mlkl^−/−^ and Mlkl^+/−^*mice (**Fig. 5G**). Given the fact that Pparγ and Pparα synergizes to induce WAT browning/beiging to enhance energy expenditure (35), we evaluated the impact of MLKL reduction or absence on browning markers [uncoupling protein 2 (Ucp2), PR/SET domain 16 (Prdm16), cell death inducing DFFA like effector a (Cidea), and Cox8b cytochrome c oxidase subunit 8B (Cox8b)] in eWAT. Absence of *Mlkl* had no effect on Ucp2 and Prdm16 levels, while Cidea and Cox8b were significantly upregulated compared to WT mice fed CD. Feeding the HFHFrHC diet significantly upregulated Ucp2, Prdm16, and Cox8b levels in *Mlkl^−/−^* mice compared to WT mice. In *Mlkl^+/−^* mice fed the HFHFrHC diet, Cidea and Cox8b were significantly upregulated compared to WT mice, while Ucp2 and Prdm16 showed a tendency for an increase without reaching statistical significance (**Fig. 5H**). Collectively, upregulation of beige adipocyte genes in combination with smaller adipocyte size distributions suggests a potential link between MLKL absence/reduction and improved insulin sensitivity in the animal, and browning of WAT contributing reduced adiposity.

### *Mlkl^−/−^* and *Mlkl^+/−^* mice exhibited improved insulin sensitivity on a HFHFrHC diet

Reduced adiposity, as well as a smaller but more adipocyte cellularity, is associated with enhanced insulin sensitivity (36, 37). We performed the glucose tolerance test (GTT) to assess the efficiency of clearance following a bolus of glucose in the WT, *Mlkl^−/−^* and *Mlkl^+/−^* groups provided CD or HFHFrHC diet. Consumption of the HFHFrHC diet led to comparable levels of glucose clearance in WT, *Mlkl^−/−^*, and *Mlkl^+/−^* mice and no difference in the area under the curve was observed (**Fig. 6A**). Next, insulin sensitivity was evaluated using the insulin tolerance test (ITT) to assess how efficiently glucose is cleared from circulation following administration of insulin. HFHFrHC feeding induced insulin resistance in WT mice, whereas *Mlkl^−/−^* and *Mlkl^+/−^*mice displayed significantly improved insulin sensitivity compared to WT mice on the same diet (**Fig. 6B**). Together, these findings suggest that loss of MLKL does not improve the body’s ability to handle a glucose load, rather, MKLK deficient mice have an increase in peripheral insulin sensitivity.

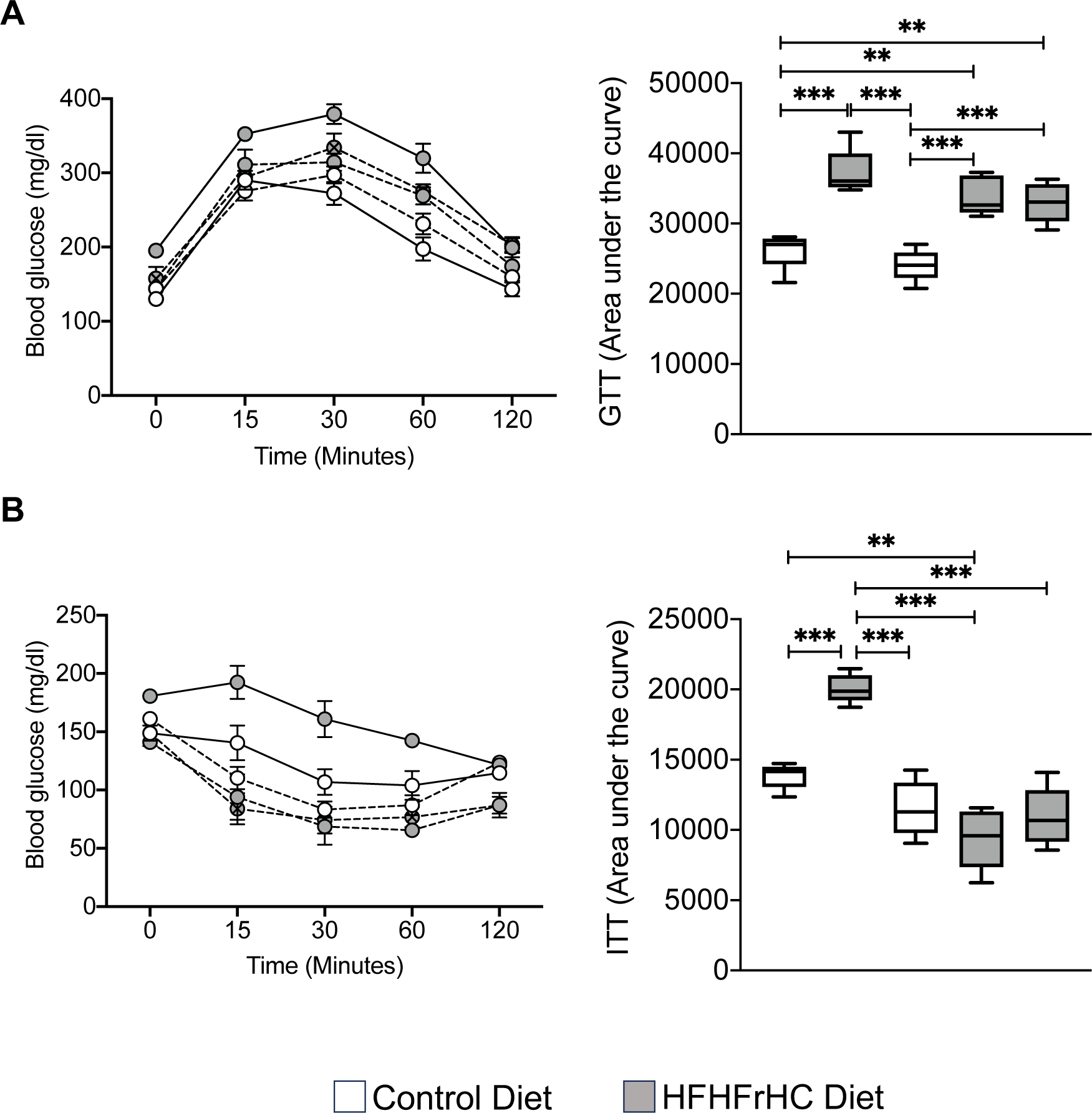

### Energy expenditure is increased in *Mlkl^−/−^* and *Mlkl^+/−^*mice on a HFHFrHC diet

We performed indirect calorimetric analysis to further evaluate the role of MLKL in whole body metabolism. The HFHFrHC diet significantly reduced oxygen consumption in WT mice when normalized to body mass during the dark phase compared to CD-fed WT mice (**Fig. 7A and Fig. S5A**). Conversely, *Mlkl^−/−^* and *Mlkl^+/−^* mice on the HFHFrHC diet exhibited increased oxygen consumption during the light (20%, 30%, respectively) and dark (50%, 50%, respectively) phases compared to WT mice on the same diet. Additionally, energy expenditure (EE) normalized to total body mass was elevated in HFHFrHC-fed *Mlkl^−/−^* and *Mlkl^+/−^* mice during both light (12%, 22%, respectively) and dark (40%, 25%, respectively) phases compared to WT mice on the same diet (**Fig. 7C, S5C**). Remarkably, when normalized to lean body mass, oxygen consumption and EE appeared comparable among HFHFrHC diet-fed WT, *Mlkl^−/−^*, and *Mlkl^+/−^*mice, suggesting a role of adipose tissue in these processes (**Figs. 7B, 7D, S5B, S5D**). Respiratory Exchange Ratio (RER) measurements, reflecting metabolic substrate utilization (38), demonstrated significantly reduced RER values (near 0.7) for HFHFrHC diet-fed groups compared to CD-fed groups, indicating that fat is used as the major substrate by WT, *Mlkl^−/−^*, and *Mlkl^+/−^* mice during both light and dark phases when the mice are fed the HFHFrHC diet (**Fig. 7E, S5E**). When comparing the RER values of HFHFrHC fed *Mlkl^−/−^* and *Mlkl^+/−^*mice in the dark phase, *Mlkl^+/−^* mice exhibited a significantly higher RER value compared to *Mlkl^−/−^* mice, indicating a potential utilization of both fat and carbohydrate sources (**Fig. 7E**).

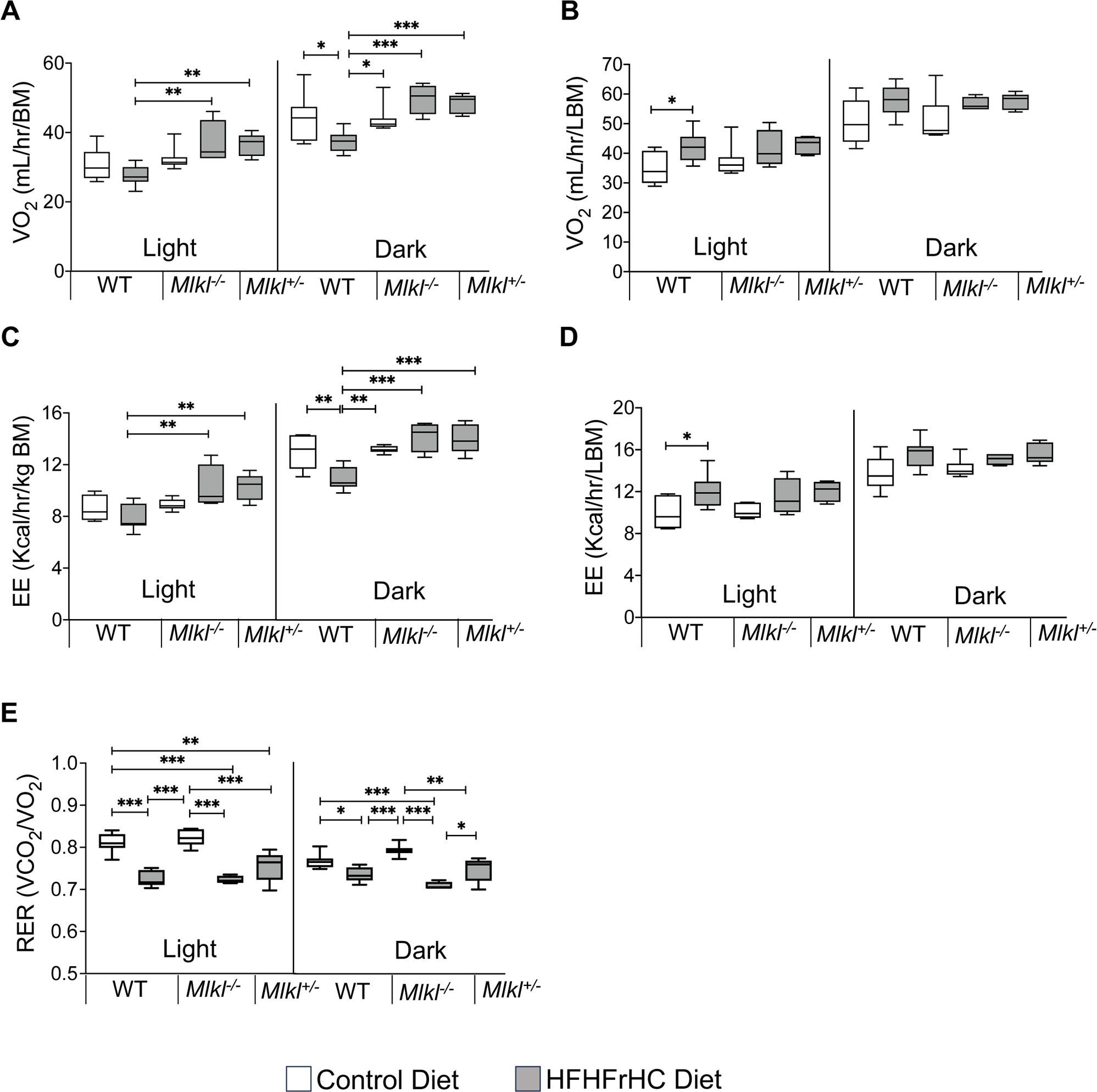

## DISCUSSION

Necroptosis, a highly proinflammatory mode of cell death, is elevated in MAFLD/MASH patients and mouse models, and inflammation is a key driver in the progression of MAFLD (12, 14, 39). Nevertheless, studies investigating the consequences of inhibiting necroptosis through targeting either RIPK3 or MLKL in mice have presented contradictory outcomes concerning inflammation, steatosis, and fibrosis, with the observed effects varying based on dietary conditions (40–43). Hence, the contribution of necroptosis to the onset and advancement of MAFLD remains uncertain. The results of our study imply that while the absence of *Mlkl* (*Mlkl^−/−^* mice) is protective against HFHFrHC diet-induced obesity and insulin resistance, it does not extend the same protection to the liver against inflammation, steatosis or fibrosis. In contrast, partial loss of *Mlkl* (*Mlkl^+/−^* mice) mitigated the adverse effects of a HFHFrHC diet on both adipose tissue and the liver, while also enhancing insulin sensitivity..

Current evidence supports a role of necroptosis and necroptosis-induced inflammation in the progression of MAFLD (18, 23, 39, 44). Contrary to these findings, we did not find an increase in necroptosis markers or necroptosis-associated proinflammatory cytokines (TNFα, IL6, IL-1β, and CCL2) in the livers of WT mice fed a HFHFrHC diet, despite observing an increase in MLKL protein. Our results align with those of Xu et al. who observed no elevation in necroptosis-related inflammatory cytokines in a MAFLD mouse model induced by a high-fat diet. Nonetheless, they did report an upsurge in necroptosis markers (22). Notably, our investigation demonstrated that the HFHFrHC diet diminishes the expression of RIPK3, a critical initiator of MLKL activation via phosphorylation. This implies a potential involvement of reduced RIPK3 expression in inhibiting necroptosis. In support of this, a recent study reported epigenetic silencing of *Ripk3* in hepatocytes of a MASH-inducing diet mouse model (45). Liver cancer cells are also shown to prevent MLKL-mediated necroptosis by epigenetic silencing of *Ripk3* (46). Surprisingly, in our study, *Mlkl^−/−^* mice on the HFHFrHC diet exhibited elevated levels of TNFα, IL6, IL1β, and Ccl2 in the liver, challenging previous data indicating that MLKL deficiency reduces inflammation (23, 44). However, this agrees with reports linking MLKL deficiency to increased proinflammatory cytokines due to the non-necroptotic function of MLKL in regulating intracellular degradation of TNF and its receptor (47). While MLKL is traditionally viewed as operating downstream of RIPK3 in necroptosis, recent reports suggest potential necroptosis-independent functions for MLKL such as autophagy, receptor internalization, ligand-receptor degradation, endosomal trafficking, extracellular vesicle generation, neutrophil trap formation, and inflammasome control (48). Moreover, the specific functions of MLKL in different cell types could also influence liver inflammation and pathology. A recent study highlighted that the absence of *Mlkl* specifically in myeloid cells exacerbated ethanol-induced hepatic injury, steatosis, and inflammation in mice (49). Therefore, further research is needed to explore the impact of cell type-specific effects of MLKL on MASH, underscoring the intricate nature of MLKL’s involvement in liver pathology and expanding its potential functions beyond conventional associations with necroptosis.

In our study, we used a HFHFrHC diet because it is widely used to induce liver fibrosis, and exhibits several clinically relevant characteristics of MASH and associated metabolic disorders in mice (24). Whereas *Mlkl^−/−^* mice did not show protection against HFHFrHC-induced liver damage, steatosis, or fibrosis, partial loss of *Mlkl* (*Mlkl^+/−^* mice) was protective. These findings are consistent with our previous data showing that absence of *Mlkl* do not protect mice against liver fibrosis induced by a choline deficient high fat diet (23). Therefore, the reported protection of mice from CCl_4_-induced or bile duct ligation-mediated liver fibrosis in the absence of *Mlkl* suggests a diet-specific influence on fibrosis distinct from the injury mediated by CCl_4_ or bile duct ligation (50, 51). The discovery that knocking out *Mlkl* specifically in endothelial cells alleviates liver fibrosis in a mouse MASH model underscores the importance of comprehending the cell-type-specific effects of MLKL in order to discern its role in MASH (52). Absence of MLKL altered lipid droplet size and distribution without affecting overall triglyceride accumulation, suggesting a potential role of MLKL in regulating lipid droplet size. The protective effect observed of the *Mlkl^+/−^* mice could be linked to increased levels of Pparα and Pparγ in the liver, which are critical regulators of liver lipid metabolism and fibrosis, respectively (53, 54). The fact that Pparα and Pparγ levels are increased only in HFHFrHC fed *Mlkl^+/−^*mice and not in WT or *Mlkl^−/−^* mice fed the same diet, hints to a dose-dependent relationship between MLKL and PPARs, necessitating further investigation into the underlying mechanisms of this novel MLKL-PPAR connection in MAFLD.

A key finding in our study is the resistance to diet-induced obesity in both *Mlkl^−/−^* and *Mlkl^+/−^* mice, consistent with similar effects reported in *Mlkl^−/−^* mice with other MAFLD-inducing diets (21, 22). Our findings show that the resistance to obesity is not due to the result of alterations in food intake or physical activity but appears to be due to an impact of *Mlkl* on adiposity. The presence of smaller adipocytes in HFHFrHC diet fed *Mlkl^−/−^* and *Mlkl^+/−^* is not due to impaired adipogenesis or *de novo* lipogenesis, rather it is due to increased fat utilization. Increased fat utilization in these mouse models is further supported by the elevated Ppar*α* levels and browning markers in white adipose tissue (35). The increased oxygen consumption and energy expenditure in both *Mlkl^−/−^* and *Mlkl^+/−^* mice is associated with fat tissue, suggest a role for MLKL in influencing fat utilization in adipose tissue. Thus, increased fat utilization by thermogenic adipocytes in the absence or reduction of MLKL contributes to the resistance to diet-induced obesity observed in these mouse models. While our data contradicts previous cell culture studies using 3T3-L1 cells, which showed that knocking down *Mlkl* impairs white adipocyte differentiation and downregulates genes involved in lipid metabolism (55), our findings *in vivo* could suggest the involvement of other factors such as adipose tissue extracellular matrix or systemic effects on adipocyte differentiation in the absence or partial loss of *Mlkl* (56–58). The fact that adipose tissue inflammation is not affected by HFHFrHC diet or MLKL supports a non-necroptotic function of MLKL in regulating lipid metabolism in adipose tissue, which warrants further investigation for potential therapeutic targets in obesity-related conditions.

The improved insulin sensitivity observed in HFHFrHC diet fed *Mlkl^−/−^*and *Mlkl^+/−^* mice, as evidenced by insulin tolerance tests, suggests a potential role for MLKL in the regulation of systemic insulin responsiveness. However, it is noteworthy that glucose tolerance remained unaltered in these mice. This observation implies that while MLKL deficiency might have specific effects on insulin-mediated glucose clearance, it might not significantly impact overall glucose homeostasis. In agreement with our results, Xu et al. (22) documented enhanced insulin sensitivity in *Mlkl^−/−^* mice on a HFD. However, diverging from our findings, they also noted improved glucose tolerance. Again, this inconsistency suggests diet as a critical factor that may contribute to the observed varied outcomes. PPARγ is a key regulator of adipocyte differentiation and function that is known to regulate systemic insulin sensitivity (26). Thus, the upregulation of Pparγ in the context of MLKL deficiency or reduction may contribute to the observed improvements in insulin sensitivity in our study.

In summary, our study underscores the diverse and non-necroptotic roles of MLKL in two key metabolic tissues, liver and white adipose tissue. Our findings challenge prior studies regarding MLKL’s involvement in liver inflammation in diet-induced MAFLD and the role of necroptosis-induced liver inflammation in fibrosis. Additionally, the study emphasizes the importance of a dose-dependent relationship of MLKL in the liver during MAFLD. Overall, the research sheds light on the intricate interplay between MLKL, lipid metabolism, and metabolic health, offering valuable insights for understanding and potentially targeting MLKL in the context of obesity-related disorders. Further, cell type-specific and mechanistic studies are warranted to elucidate the precise pathways through which MLKL exerts its influence on these physiological processes. The translational implications of these findings may offer new avenues for therapeutic interventions in conditions characterized by metabolic dysfunction and liver pathology.

## MATERIALS AND METHODS

### Animals and diet feeding

All procedures were approved by the Institutional Animal Care and Use Committee at the University of Oklahoma Health Sciences Center (OUHSC). *Mlkl^−/−^* mice were developed by Murphy et al (2013) (59) and in our study, colonies of WT (*Mlkl^+/+^*)*, Mlkl^−/−^* and *Mlkl^+/−^* mice were generated by breeding male and female *Mlkl^+/−^* mice. The WT (*Mlkl^+/+^*) mice used in the study are littermates from *Mlkl^+/−^* mice cross-breeding. The mice were group housed in ventilated cages at 20 ± 2 °C, 12-h/12-h dark/light cycle. Diet feeding was done for 6 months, starting at 2 months of age. Male mice (n=5-8/group) were fed either a high-fat, high-fructose, high cholesterol diet (HFHFrHC) containing 40 kcal% Fat (Mostly Palm Oil), 20 kcal% Fructose, and 2% Cholesterol (D09100310, Research Diets Inc, New Brunswick, NJ) or a matched control diet (D09100304,Research Diets Inc, New Brunswick, NJ) *ad libitum* for a period of 6 months at the OUHSC animal care facility. Detailed diet composition is presented in supplementary **Table S3** and experimental design is presented in **Fig. 1A**.

### Glucose tolerance and insulin tolerance tests (GTT and ITT)

GTT and ITT were performed as described before (60). In brief, mice were fasted for 6 hours (for GTT) or 5 hours (for ITT) and received an intraperitoneal injection of glucose (1g/kg, Sigma Aldrich) or insulin (0.75 units/kg, Novo Nordisk Inc., Denmark). Glucose concentration was determined from blood samples collected from tail vein after 15, 30, 60and 120 minutes of glucose or insulin challenge using TRUE METRIX glucose strips and glucometer (Trividia Health Inc., USA).

### Histological analysis of liver sections

Paraffin embedded liver or visceral fat sections were stained with Hematoxylin & Eosin (H&E) according to the standardized protocol at the Stephenson Cancer Center Tissue Pathology core. Images were acquired using a Nikon Ti Eclipse microscope (Nikon, Melville, NY) at 20x magnification for 3 random non overlapping fields per sample. Lipid droplet number and area were quantified using Adiposoft Fiji extension software as described (61) and are represented graphically.

### Picrosirius red staining

Picrosirius red staining was performed with paraffin embedded liver sections (4 µM) by following a standardized protocol at the Imaging Core facility at the Stephenson Cancer Center Tissue Pathology core. Images were acquired using Nikon Ti Eclipse microscope (Nikon, Melville, NY) for 3 random non-overlapping fields per sample at 20X magnification and quantified using Image J software (U.S. National Institutes of Health).

### Alanine transaminase (ALT) and Triglyceride Assessment

Plasma ALT levels, plasma and hepatic triglycerides were quantified using Alanine transaminase or Triglyceride colorimetric assay kits obtained from Cayman Chemical Company (Ann Arbor, MI). The measurements were conducted in accordance with the manufacturer’s instructions.

### Hydroxyproline assay

The collagen content in the liver was measured by hydroxyproline content as described (23). The absorbance values at 558nm were converted into µg units using the 4-parameter standard curve generated using the standards and expressed as µg hydroxyproline/g of tissue.

### Indirect Calorimetry studies

Indirect calorimetry was performed using a 16-chamber metabolic monitoring system (Syble Systems Itnl., Promethion) for the continuous monitoring of CO_2_ production and O_2_ consumption in individual mice. Mice destined for metabolic monitoring were acclimated to the system for 48-72 hours prior to data collection periods. Following acclimation, mice were monitored continuously for 72 hours, during which time energy intake (EI) was measured in real time. Energy intake, respiratory exchange ratio (vCO_2_/vO_2_), energy expenditure and energy balance measurements were calculated as an average of measures over the period of study. Activity was determined by infrared beam breaks as a measure of horizontal and vertical movement (XYZ-axis) of mice. The food and water intake amount and the frequency/duration of feeding behavior within the 24–120-hour experimental window was quantified by the Promethion system. Body composition was quantified by magnetic resonance with a 1 min primary accumulation time (4-in-1 700 EchoMRI, Houston, TX).

### Western Blot Analysis

Western blot analysis was performed and quantified using ImageJ software (U.S. National Institutes of Health, Bethesda, MD, USA) as described previously (18). The primary antibodies used were: MLKL (MABC60, Millipore); P-MLKL (#ab196436, Abcam); RIPK3 (#NBP1-77299, 117 Novus Biologicals), β-tubulin (sc-5274, Santa Cruz Biotechnology). In the representative western blots, each experimental group is represented by 4 independent mice. Graphical representation of the quantified western blots for 4 animals/group are shown alongside the blots.

### Quantitative real-time PCR (RT-PCR)

Briefly, total RNA was isolated using RNeasy kit (Qiagen, Valencia, CA), first-strand cDNA was synthesized using High-capacity cDNA reverse transcription kit (Thermo Fisher Scientific, Waltham, MA), and. RT-PCR using Power SYBR Green PCR Master Mix (Thermo Fisher Scientific, Waltham, MA) in a Quantstudio 12K Flex real time PCR system (Applied Biosystems). Primers used for RT-PCR are listed in **Table S1**. PCR arrays were performed using RT^2^ Profiler™ PCR Array Mouse Cytokines & Chemokines (PAMM-150Z, Qiagen). Calculations were performed by a comparative method (2^−ΔΔ^Ct) using β-microglobulin, β**-**actin, or hypoxanthine phosphoribosyltransferase 1 (HPRT) as controls, as described previously (23). The expression of genes, comparison of signatures between the groups and heat map representation was performed using Microsoft Excel (Version 16.78.3, 2023) and MetaboAnalyst 5.0.

### Quantitative Targeted Proteomics

Quantitative proteomics was used to determine changes in mitochondrial enzymes in liver tissue as previously described (18, 62). Briefly, 20 μg total liver tissue homogenate was run 1.5 cm into a 12.5% SDS–PAGE gel (Criterion, Bio-Rad). This was followed by fixation and staining with GelCode Blue (Pierce). The entire lane was cut into ∼1 mm3 pieces, washed, reduced with DTT, alkylated with iodoacetamide, and digested with trypsin. The peptides generated were extracted with 50% methanol/10% formic acid in water, dried, reconstituted in 1% acetic acid, and analyzed using selected reaction monitoring (SRM) with a triple quadrupole mass spectrometer (ThermoScientific TSQ Quantiva) configured with a splitless capillary column HPLC system (ThermoScientific Ultimate 3000). Data processing was done using the program Skyline, which aligned the various collision-induced dissociation reactions monitored for each peptide and determines the chromatographic peak areas. The response for each protein was taken as the total response for all peptides monitored. Changes in the relative abundance of the proteins were determined by normalization to the BSA internal standard, with confirmation by normalization to the housekeeping proteins.

### Statistical Analysis

All quantitative data are represented as mean ±SEM. One-way ANOVA with Tukey’s multiple comparison’s test was used to analyze data using GraphPad Prism (Version 10.0.3). P < 0.05 is considered statistically significant. The symbols used for statistical comparison between groups are described in the figure legend.

## ACKNOWLEDGEMENTS

The efforts of authors were supported by NIH grants R01AG059718 (S.S. Deepa), R03 CA262044 (S.S. Deepa), Oklahoma Center for Adult Stem Cell Research Grant (OCASCR; S.S. Deepa), Oklahoma Center for the Advancement of Science and Technology (OCAST) Postdoctoral grant HF21-009 (S. Mohammed), Department of Veterans Affairs: Senior Career Research Award I01BX004538 (A. Richardson), Presbyterian Health Foundation award under the PHF Equipment Grant award mechanism (M. Rudolph), NIH grants 5R24GM137786, 5P30AG050911, and 5P20GM103447 (M. Kinter). The authors would like to thank Stephenson Cancer Center Tissue Pathology Core for performing H&E staining, the Imaging Core facility at the Oklahoma Medical Research Foundation for performing Picrosirius red staining.

## CONFLICT OF INTERST

None.

## SUPPLEMENTARY FIGURE LEGENDS

**Figure S1:**
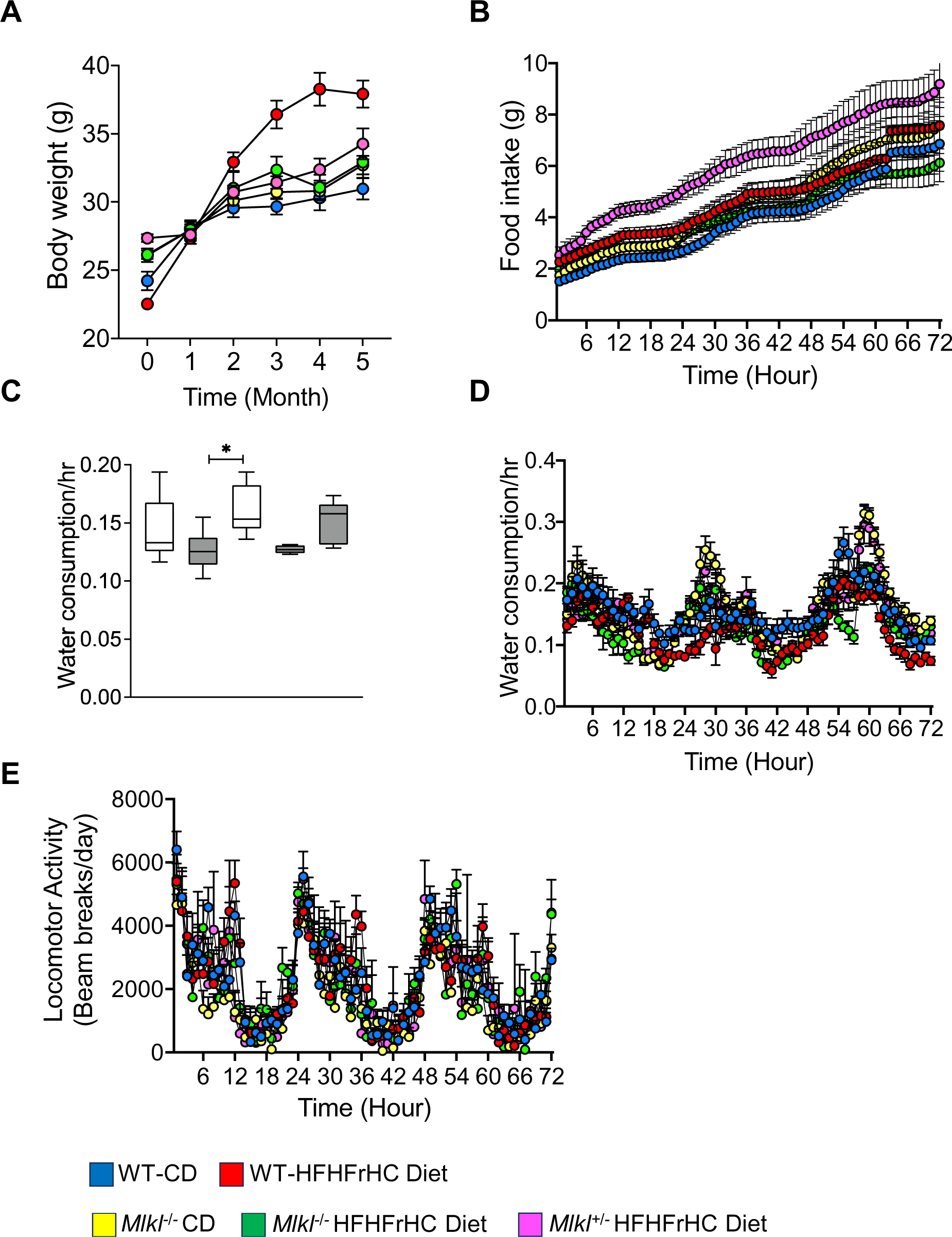
Changes in body weight, food intake, water consumption, and locomotor activity of CD or HFHFrHC diet fed WT, *Mlkl^−/−^* or *Mlkl^+/−^* mice. **(A)** Change in body weight during diet feeding from baseline through six months; Indirect calorimetry measurements over a period of 72 hours after 6 months of diet feeding showing daily food intake **(B)**, average water consumption **(C)**, daily profile of water consumption **(D)**, and daily profile of spontaneous locomotor activity **(E)**. In the figure-Blue: WT mice on CD; Red: WT mice on HFHFrHC diet; Yellow: *Mlkl*^−/−^ mice on CD; Green: *Mlkl*^−/−^ mice on HFHFrHC diet and Pink: *Mlkl*^+/−^ mice on HFHFrHC diet. (n= 7-8 for WT; 5-8 for *Mlkl*^−/−^ or *Mlkl*^+/−^). White and gray bars box plots represent experimental groups fed either CD or HFHFrHC diet, respectively. Error bars are represented as mean± SEM. One-way ANOVA P0.05, * p< 0.05, ** p< 0.005, ***p< 0.0005

**Figure S2:**
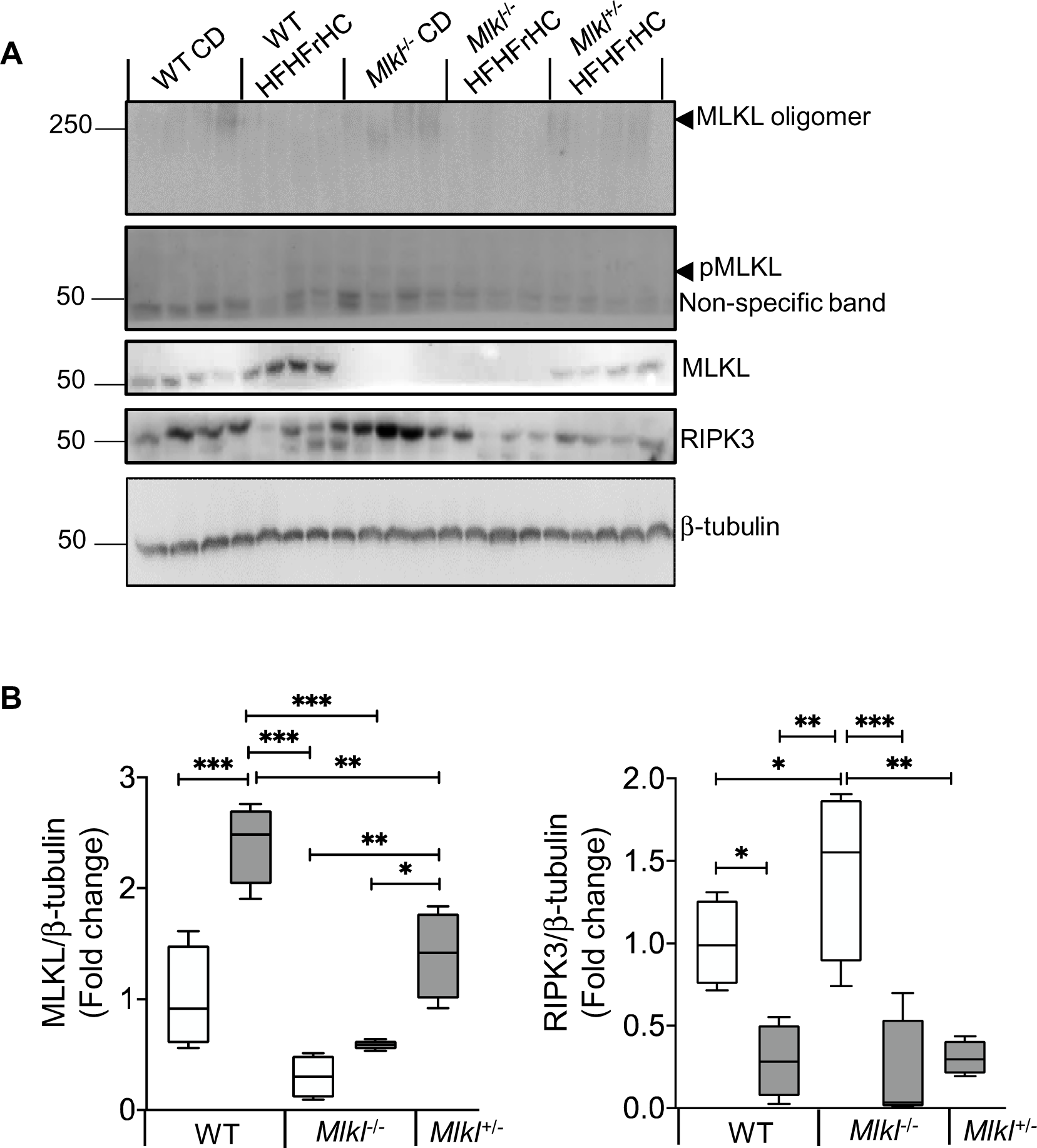
Necroptosis markers in the livers of CD or HFHFrHC diet fed mice. **(A)** Immunoblots of liver tissue extracts for necroptosis proteins: RIPK3, MLKL, pMLKL, MLKL oligomer, and loading control β-tubulin from WT, *Mlkl^−/−^* or *Mlkl^+/−^* fed test diets; **(B)** Graphical representation of quantified blot normalized to β-tubulin. White and gray bars box plots represent experimental groups fed either CD or HFHFrHC, respectively. Error bars are represented as mean± SEM. One-way ANOVA P0.05, * p< 0.05, ** p< 0.005, ***p< 0.0005

**Figure S3:**
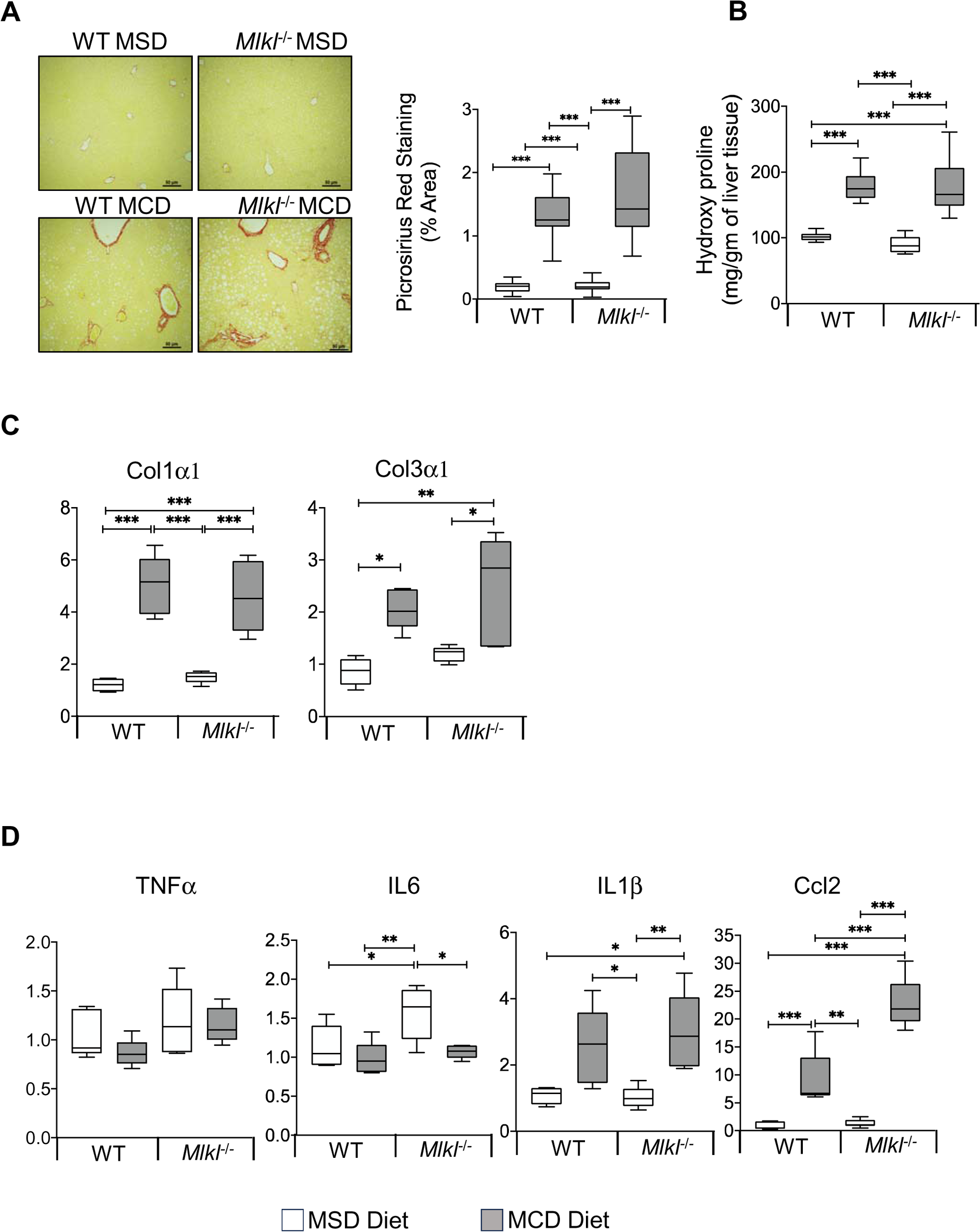
Fibrosis markers in the livers of MSD or MCD diet fed mice. **(A)** *Left:* Representative PSR staining of liver sections of WT, *Mlkl^−/−^* fed either methionine-choline sufficient diet (MSD) or methionine-choline deficient (MCD) diet. Scale bar: 50 mM. *Right:* Graphical representation of PSR quantification. (n=3/group); **(B)** Estimation of total hydroxyproline content in liver tissues of WT, *Mlkl^−/−^* mice fed MSD or MCD (n=5/group); **(C)** The transcript levels fibrosis markers Col1a1 and Col3a1 and inflammatory cytokine markers TNFα, IL6, IL1β, and Ccl2 **(D)** in the livers of WT, *Mlkl^−/−^* fed MSD, or MCD. Each transcript is represented as a fold change normalized to β-microglobulin (n=6/group). White and gray bars box plots represent experimental groups fed either CD or HFHFrHC, respectively. Error bars are represented as mean± SEM. One-way ANOVA P0.05, * p< 0.05, ** p< 0.005, ***p< 0.0005.

**Figure S4:**
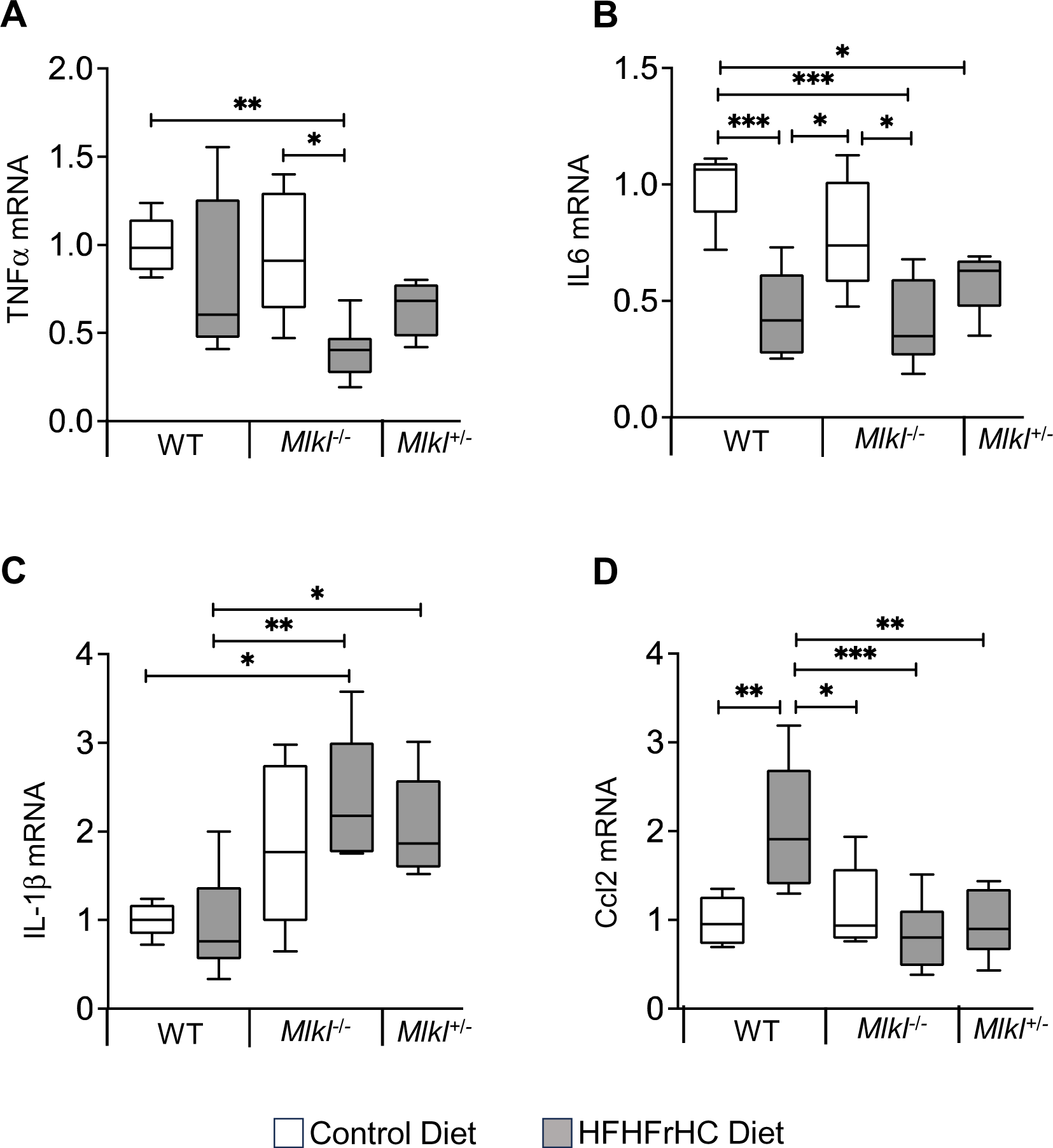
Effect of absence or reduction of *Mlkl* on eWAT inflammatory cytokines. Data from WT, *Mlkl^−/−^* or *Mlkl^+/−^* mice fed a CD (white box plots) or HFHFrHC diet (box plots): Transcript levels of inflammatory cytokines TNFα **(A)**, IL6 **(B)**, IL1β **(C)**, and Ccl2 **(D)**, normalized to β-microglobulin and represented as fold change relative to CD fed WT mice (n= 7-8 for WT; 5-8 for *Mlkl*^−/−^ or *Mlkl*^+/−^). Error bars are represented as mean± SEM. One-way ANOVA P0.05, * p< 0.05, ** p< 0.005, ***p< 0.0005.

**Figure S5:**
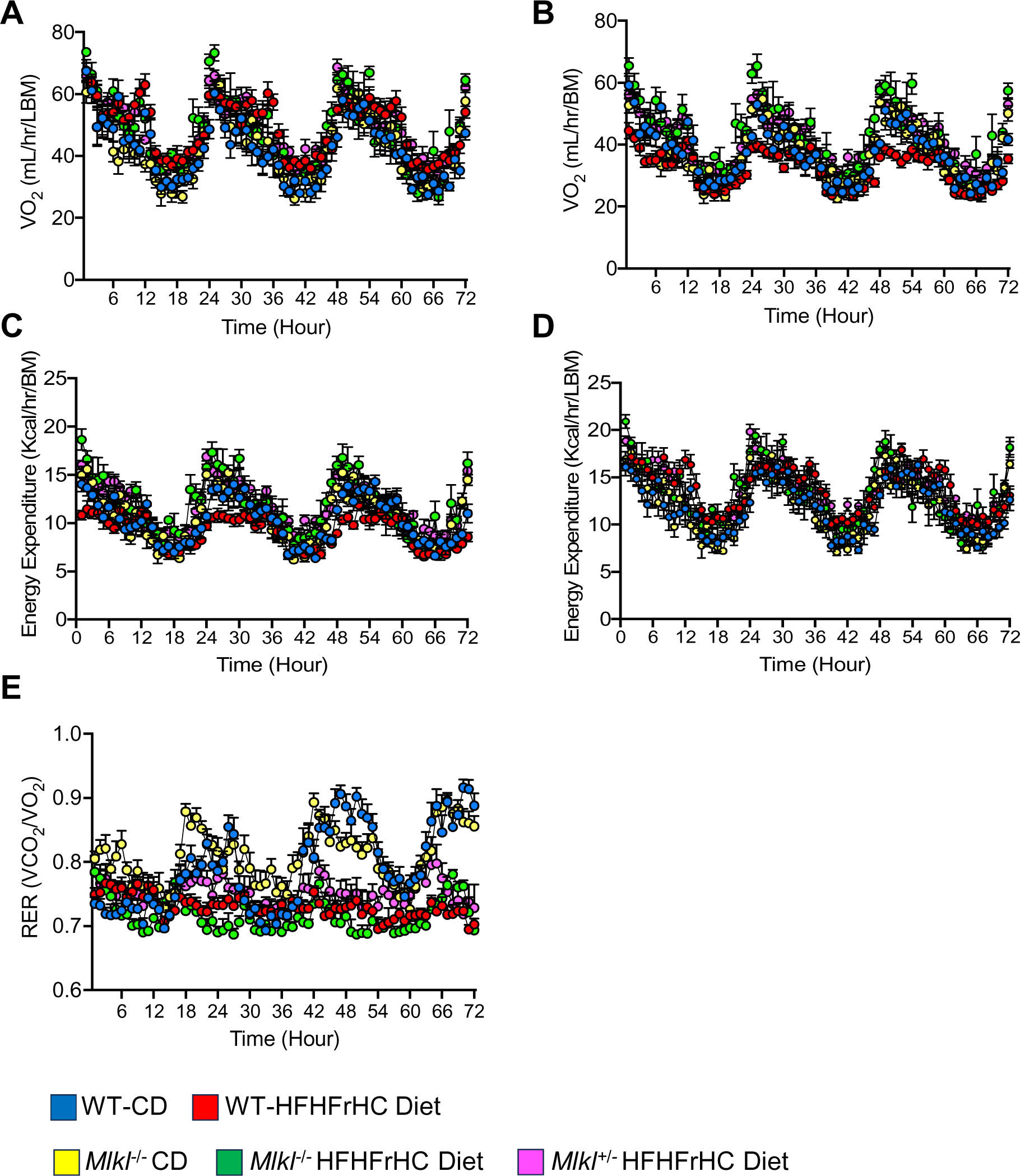
Time course changes in oxygen consumption rate, energy expenditure, and respiratory quotient ratio of WT, *Mlkl^−/−^* or *Mlkl^+/−^* fed test diets. Measures were recorded by indirect calorimetry over 72 hours in 6-hour intervals. Daily profiles of oxygen consumption rate normalized to total body mass (BM) **(A)** and lean body mass (LBM) **(B)**; Daily profiles of energy expenditure normalized to total body mass (BM) **(C)** and lean body mass (LBM) **(D)**; **(E)** Daily profile of respiratory quotient. In the figure legend-Blue: WT mice on LFD; Red: WT mice on HFD; Yellow: *Mlkl*^−/−^ mice on LFD; Green: *Mlkl*^−/−^ mice on HFD and Pink: *Mlkl*^+/−^ mice on HFD. (n= 7-8 for WT; 5-8 for *Mlkl*^−/−^ or *Mlkl*^+/−^).

## Notes

### Competing Interest Statement

The authors have declared no competing interest.

