## Supplementary material for "Non-Necroptotic Roles of MLKL in Diet-Induced Obesity, Liver Pathology, and Insulin Sensitivity: Insights from a High Fat, High Fructose, High Cholesterol Diet Mouse Model": Table 1

**Table S1**

| **Gene** | **Forward Sequence** | **Reverse Sequence** |
| --- | --- | --- |
| β-microglobulin | 5′-CACTGACCGGCCTGTATGC-3′ | 5′-GGGTGGCGTGAGTATACTTGAAT-3′ |
| Ccl2 | 5’-TTAAAAACCTGGATCGGAACCAA-3’ | 5’-GCATTAGCTTCAGATTTACGGGT-3’ |
| Cebpα | 5’-GCAAAGCCAAGAAGTCGGTGGA-3’ | 5’-CCTTCTGTTGCGTCTCCACGTT-3’ |
| Cidea | 5’-GGTGGACACAGAGGAGTTCTTTC-3’ | 5’-CGAAGGTGACTCTGGCTATTCC-3’ |
| Cox8b | 5’-GCGAAGTTCACAGTGGTTCC-3’ | 5’-GGAACCATGAAGCCAACGAC-3’ |
| Hprt | 5’-CTGGTGAAAAGGACCTCTCG-3’ | 5’-TGAAGTACTCATTATAGTCAAGGGCA-3’ |
| IL-6 | 5’-TGGTACTCCAGAAGACCAGAGG-3’ | 5’-AACGATGATGCACTTGCAGA-3’ |
| Il-1b | 5’-AGGTCAAAGGTTTGGAAGCA-3’ | 5’-TGAAGCAGCTATGGCAACTG-3’ |
| TNFα | 5’-CACAGAAAGCATGATCCGCGACGT-3’ | 5’- CGGCAGAGAGGAGGTTGACTTTCT-3’ |
| Pparα | 5’-ACCACTACGGAGTTCACGCATG-3’ | 5′-GAATCTTGCAGCTCCGATCACAC-3’ |
| Pparγ | 5’-GTACTGTCGGTTTCAGAAGTGCC-3’ | 5′-ATCTCCGCCAACAGCTTCTCCT-3’ |
| Prdm16 | 5′-AGCTGCGATTGAGGAAGAGCGA-3′ | 5′-CCTGGGTTCTGACGCCTTTGTT-3′ |
| Ucp2 | 5′-CGTTCTGGGTACCATCCTAACC-3′ | 5′–CAAGCCTCCATCCAAGTGTCAC–3′ |
