## Supplementary material for "Non-Necroptotic Roles of MLKL in Diet-Induced Obesity, Liver Pathology, and Insulin Sensitivity: Insights from a High Fat, High Fructose, High Cholesterol Diet Mouse Model": Table 2

| Genes | WT-LFD | | WT-HFHFrHC | | Mlkl^-/-^LFD | | Mlkl^-/-^HFHFrHC | | Mlkl^+/-^HFHFrHC | | Significant differences between the groups | | | | |
| --- | --- | --- | --- | --- | --- | --- | --- | --- | --- | --- | --- | --- | --- | --- | --- |
|  | Fold Change | SEM | Fold Change | SEM | Fold Change | SEM | Fold Change | SEM | Fold Change | SEM | WT-LFD vs. WT-HFHFrHC | WT-LFD vs. Mlkl^-/-^LFD | WT-HFHFrHC vs. Mlkl^-/-^HFHFrHC | WT-HFHFrHC vs. Mlkl^+/-^HFHFrHC | Mlkl^-/-^HFHFrHC vs. MLKL^+/-^HFHFrHC |
| Adipoq | 1 | 0.18 | 0.14 | 0.03 | 0.38 | 0.15 | 1.17 | 0.26 | 1.43 | 0.33 | No | No | Yes | Yes | No |
| Bmp2 | 1 | 0.28 | 0.71 | 0.08 | 1.18 | 0.16 | 1.29 | 0.36 | 0.94 | 0.16 | No | No | No | No | No |
| Bmp4 | 1 | 0.23 | 1.51 | 0.24 | 2.71 | 0.07 | 6.94 | 0.29 | 3.94 | 0.69 | No | No | Yes | Yes | Yes |
| Bmp6 | 1 | 0.19 | 0.51 | 0.10 | 1.52 | 0.16 | 1.77 | 0.41 | 1.26 | 0.24 | No | No | Yes | No | No |
| Bmp7 | 1 | 0.24 | 1.81 | 0.38 | 1.90 | 0.27 | 2.52 | 0.22 | 3.77 | 0.33 | No | No | No | Yes | No |
| Ccl1 | 1 | 0.24 | 1.99 | 0.90 | 0.87 | 0.19 | 10.93 | 2.99 | 4.46 | 2.61 | No | No | Yes | No | No |
| Ccl11 | 1 | 0.20 | 2.79 | 0.49 | 1.67 | 0.13 | 32.96 | 2.37 | 5.56 | 1.25 | No | No | Yes | No | Yes |
| Ccl12 | 1 | 0.24 | 4.87 | 0.89 | 1.02 | 0.12 | 13.00 | 5.31 | 3.83 | 2.23 | No | No | No | No | No |
| Ccl17 | 1 | 0.27 | 4.10 | 1.64 | 0.87 | 0.20 | 1.61 | 0.67 | 1.04 | 0.49 | No | No | No | No | No |
| Ccl19 | 1 | 0.12 | 2.03 | 0.26 | 1.01 | 0.07 | 5.08 | 0.38 | 4.94 | 2.00 | No | No | No | No | No |
| Ccl2 | 1 | 0.41 | 6.25 | 1.85 | 2.89 | 1.67 | 15.26 | 4.84 | 3.48 | 1.49 | No | No | No | No | Yes |
| Ccl20 | 1 | 0.26 | 2.57 | 1.12 | 0.68 | 0.10 | 14.13 | 2.92 | 1.78 | 1.04 | No | No | Yes | No | No |
| Ccl22 | 1 | 0.33 | 42.07 | 0.47 | 3.71 | 0.76 | 46.30 | 8.68 | 96.27 | 18.10 | Yes | No | No | Yes | Yes |
| Ccl24 | 1 | 0.27 | 1.74 | 0.85 | 1.22 | 0.08 | 5.65 | 1.21 | 4.63 | 1.98 | No | No | No | No | No |
| Ccl3 | 1 | 0.26 | 5.52 | 2.77 | 1.06 | 0.25 | 13.87 | 3.34 | 11.91 | 3.48 | No | No | No | No | No |
| Ccl4 | 1 | 0.21 | 4.81 | 1.95 | 1.05 | 0.16 | 12.87 | 2.22 | 12.45 | 2.97 | No | No | No | No | No |
| Ccl5 | 1 | 0.10 | 2.60 | 0.29 | 1.30 | 0.41 | 6.64 | 0.82 | 5.82 | 2.60 | No | No | No | No | No |
| Ccl7 | 1 | 0.57 | 6.99 | 2.94 | 5.14 | 0.05 | 4.74 | 3.51 | 0.89 | 0.13 | No | No | No | No | No |
| Cd40lg | 1 | 0.12 | 0.31 | 0.11 | 0.70 | 0.14 | 2.66 | 0.51 | 1.97 | 0.10 | No | No | Yes | Yes | No |
| Cd70 | 1 | 0.24 | 0.70 | 0.09 | 0.87 | 0.19 | 1.63 | 0.40 | 1.75 | 0.21 | No | No | No | Yes | No |
| Cntf | 1 | 0.19 | 0.58 | 0.12 | 0.85 | 0.17 | 1.23 | 0.08 | 1.33 | 0.21 | No | No | No | Yes | No |
| Csf1 | 1 | 0.16 | 0.89 | 0.14 | 0.84 | 0.19 | 2.27 | 0.35 | 2.05 | 0.70 | No | No | No | No | No |
| Csf2 | 1 | 0.24 | 0.28 | 0.06 | 0.32 | 0.11 | 4.53 | 0.76 | 4.65 | 0.39 | No | No | Yes | Yes | No |
| Csf3 | 1 | 0.09 | 0.98 | 0.04 | 2.40 | 0.43 | 1.68 | 0.42 | 13.45 | 2.89 | No | No | No | Yes | Yes |
| Ctf1 | 1 | 0.18 | 0.92 | 0.21 | 1.42 | 0.22 | 1.95 | 0.37 | 1.99 | 0.53 | No | No | No | No | No |
| Cx3cl1 | 1 | 0.18 | 2.58 | 1.27 | 1.95 | 0.25 | 13.15 | 3.80 | 4.99 | 1.72 | No | No | Yes | No | Yes |
| Cxcl1 | 1 | 0.64 | 0.86 | 0.14 | 0.58 | 0.26 | 1.38 | 0.46 | 9.25 | 3.49 | No | No | No | Yes | Yes |
| Cxcl10 | 1 | 0.73 | 1.60 | 0.17 | 1.28 | 0.26 | 2.77 | 0.47 | 1.93 | 0.64 | No | No | No | No | No |
| Cxcl11 | 1 | 0.20 | 0.34 | 0.08 | 0.79 | 0.19 | 1.02 | 0.23 | 0.85 | 0.42 | No | No | No | No | No |
| Cxcl12 | 1 | 0.22 | 0.96 | 0.06 | 1.46 | 0.22 | 2.17 | 0.22 | 1.98 | 0.03 | No | No | Yes | Yes | No |
| Cxcl13 | 1 | 0.20 | 0.19 | 0.05 | 1.37 | 0.19 | 1.01 | 0.54 | 1.04 | 0.33 | No | No | No | No | No |
| Cxcl16 | 1 | 0.16 | 2.14 | 0.54 | 1.23 | 0.17 | 7.14 | 1.72 | 1.60 | 0.47 | No | No | Yes | No | Yes |
| Cxcl3 | 1 | 0.24 | 0.76 | 0.03 | 0.81 | 0.48 | 2.70 | 1.11 | 0.75 | 0.28 | No | No | No | No | No |
| Cxcl5 | 1 | 0.24 | 0.76 | 0.03 | 1.04 | 0.22 | 17.12 | 6.24 | 1.75 | 0.21 | No | No | Yes | No | Yes |
| Cxcl9 | 1 | 0.07 | 3.91 | 0.66 | 1.57 | 0.29 | 9.35 | 1.84 | 7.86 | 0.85 | No | No | Yes | No | No |
| Fasl | 1 | 0.25 | 1.44 | 0.43 | 0.85 | 0.18 | 4.90 | 1.89 | 0.41 | 0.05 | No | No | No | No | Yes |
| Gpi1 | 1 | 0.16 | 0.50 | 0.09 | 0.73 | 0.13 | 1.38 | 0.33 | 1.01 | 0.28 | No | No | No | No | No |
| Hc | 1 | 0.32 | 0.91 | 0.19 | 1.69 | 0.20 | 2.23 | 0.54 | 2.01 | 0.24 | No | No | Yes | No | No |
| Ifna2 | 1 | 0.40 | 12.65 | 3.70 | 1.34 | 0.12 | 4.58 | 0.67 | 15.04 | 1.96 | Yes | No | No | No | Yes |
| Ifng | 1 | 0.53 | 2.43 | 0.25 | 1.24 | 0.12 | 10.96 | 2.06 | 0.76 | 0.11 | No | No | Yes | No | Yes |
| Il10 | 1 | 0.24 | 4.18 | 1.03 | 1.16 | 0.17 | 5.06 | 0.66 | 11.32 | 3.37 | No | No | No | Yes | No |
| Il11 | 1 | 0.24 | 5.28 | 0.89 | 3.98 | 0.94 | 3.80 | 0.67 | 1.75 | 0.21 | Yes | Yes | No | Yes | No |
| Il12a | 1 | 0.24 | 1.69 | 0.09 | 2.16 | 0.05 | 3.64 | 0.27 | 1.75 | 0.21 | No | Yes | Yes | No | Yes |
| Il12b | 1 | 0.24 | 15.35 | 3.98 | 2.65 | 0.37 | 35.37 | 3.71 | 20.69 | 4.08 | Yes | No | Yes | No | Yes |
| Il13 | 1 | 0.24 | 1.43 | 0.56 | 5.87 | 1.13 | 7.08 | 1.66 | 1.75 | 0.21 | No | Yes | Yes | No | Yes |
| Il15 | 1 | 0.33 | 1.43 | 0.40 | 1.18 | 0.11 | 4.77 | 0.71 | 2.93 | 0.56 | No | No | Yes | No | No |
| Il16 | 1 | 0.29 | 1.04 | 0.16 | 1.10 | 0.27 | 3.15 | 0.45 | 2.61 | 0.61 | No | No | Yes | No | No |
| Il17a | 1 | 0.24 | 0.76 | 0.03 | 1.02 | 0.12 | 3.55 | 0.54 | 1.75 | 0.21 | No | No | Yes | Yes | Yes |
| Il17f | 1 | 0.14 | 0.86 | 0.09 | 1.56 | 0.11 | 2.82 | 1.01 | 6.54 | 3.60 | No | No | No | No | No |
| Il18 | 1 | 0.32 | 0.65 | 0.19 | 1.76 | 0.30 | 1.85 | 0.39 | 1.51 | 0.40 | No | No | No | No | No |
| Il1a | 1 | 0.21 | 1.00 | 0.20 | 1.38 | 0.35 | 2.94 | 0.77 | 3.11 | 1.32 | No | No | No | No | No |
| Il1b | 1 | 0.22 | 0.93 | 0.26 | 1.48 | 0.03 | 4.68 | 1.01 | 1.81 | 0.70 | No | No | Yes | No | Yes |
| Il1rn | 1 | 0.31 | 16.20 | 3.10 | 1.74 | 0.38 | 21.29 | 3.26 | 9.40 | 1.57 | Yes | No | No | No | Yes |
| Il2 | 1 | 0.24 | 1.36 | 0.45 | 0.87 | 0.19 | 7.58 | 3.98 | 1.75 | 0.21 | No | No | Yes | No | Yes |
| Il21 | 1 | 0.24 | 0.76 | 0.03 | 0.87 | 0.19 | 3.47 | 0.87 | 1.75 | 0.21 | No | No | Yes | No | Yes |
| Il22 | 1 | 0.24 | 0.76 | 0.03 | 1.01 | 0.10 | 1.11 | 0.15 | 1.75 | 0.21 | No | No | No | Yes | No |
| Il23a | 1 | 0.28 | 0.67 | 0.17 | 0.52 | 0.16 | 1.35 | 0.46 | 1.36 | 0.29 | No | No | No | No | No |
| Il24 | 1 | 0.24 | 0.76 | 0.03 | 0.87 | 0.19 | 0.54 | 0.29 | 1.75 | 0.21 | No | No | No | Yes | Yes |
| Il27 | 1 | 0.12 | 2.79 | 0.33 | 1.44 | 0.14 | 4.89 | 0.18 | 5.41 | 0.89 | Yes | No | Yes | Yes | No |
| Il3 | 1 | 0.24 | 0.76 | 0.03 | 1.26 | 0.30 | 2.36 | 0.06 | 1.75 | 0.21 | No | No | Yes | Yes | No |
| Il4 | 1 | 0.28 | 1.73 | 0.35 | 2.78 | 1.02 | 4.12 | 1.48 | 2.43 | 1.89 | No | No | No | No | No |
| Il5 | 1 | 0.24 | 4.49 | 1.17 | 2.28 | 0.10 | 3.01 | 0.58 | 1.75 | 0.21 | Yes | No | No | No | No |
| Il6 | 1 | 0.24 | 2.29 | 0.34 | 2.48 | 0.31 | 5.70 | 1.05 | 1.75 | 0.21 | No | No | Yes | No | Yes |
| Il7 | 1 | 0.38 | 1.39 | 0.27 | 0.60 | 0.01 | 6.29 | 0.71 | 2.98 | 0.70 | No | No | Yes | No | Yes |
| Il9 | 1 | 0.24 | 2.32 | 0.52 | 0.49 | 0.24 | 1.11 | 0.13 | 1.75 | 0.21 | Yes | No | No | No | No |
| Lif | 1 | 0.24 | 5.47 | 1.86 | 4.12 | 0.17 | 35.60 | 4.41 | 1.75 | 0.21 | No | No | Yes | No | Yes |
| Lta | 1 | 0.24 | 1.97 | 0.60 | 0.87 | 0.19 | 11.68 | 4.69 | 10.18 | 4.51 | No | No | No | No | No |
| Ltb | 1 | 0.33 | 2.62 | 0.90 | 1.36 | 0.39 | 10.61 | 3.03 | 5.18 | 0.83 | No | No | Yes | No | Yes |
| Mif | 1 | 0.11 | 1.11 | 0.18 | 1.41 | 0.27 | 1.47 | 0.19 | 1.12 | 0.10 | No | No | No | No | No |
| Mstn | 1 | 0.24 | 0.76 | 0.03 | 0.53 | 0.22 | 3.69 | 2.11 | 1.75 | 0.21 | No | No | No | No | No |
| Nodal | 1 | 0.16 | 0.49 | 0.11 | 0.43 | 0.06 | 2.23 | 0.53 | 2.49 | 1.09 | No | No | No | No | No |
| Osm | 1 | 0.24 | 4.90 | 1.01 | 1.81 | 0.33 | 16.98 | 4.20 | 1.75 | 0.21 | No | No | Yes | No | Yes |
| Pf4 | 1 | 0.21 | 2.11 | 0.81 | 0.84 | 0.13 | 5.34 | 0.99 | 1.87 | 0.87 | No | No | Yes | No | Yes |
| Ppbp | 1 | 0.28 | 1.41 | 0.31 | 0.91 | 0.19 | 2.57 | 0.72 | 3.65 | 1.30 | No | No | No | No | No |
| Spp1 | 1 | 0.09 | 6.09 | 2.01 | 1.71 | 0.45 | 53.54 | 6.57 | 2.94 | 1.58 | No | No | Yes | No | Yes |
| Tgfb2 | 1 | 0.18 | 1.46 | 0.44 | 0.50 | 0.04 | 3.56 | 0.84 | 1.33 | 0.37 | No | No | Yes | No | Yes |
| Thpo | 1 | 0.06 | 0.57 | 0.04 | 1.08 | 0.09 | 1.58 | 0.08 | 1.22 | 0.48 | No | No | No | No | No |
| Tnf | 1 | 0.52 | 3.30 | 0.82 | 1.29 | 0.52 | 11.28 | 4.10 | 6.64 | 2.18 | No | No | Yes | No | No |
| Tnfrsf11b | 1 | 0.19 | 1.90 | 0.24 | 1.18 | 0.26 | 2.43 | 0.87 | 1.76 | 0.36 | No | No | No | No | No |
| Tnfsf10 | 1 | 0.55 | 1.04 | 0.44 | 1.46 | 0.49 | 3.20 | 0.36 | 2.23 | 0.69 | No | No | Yes | No | No |
| Tnfsf11 | 1 | 0.26 | 0.76 | 0.03 | 0.73 | 0.14 | 0.98 | 0.06 | 1.51 | 0.18 | No | No | No | Yes | No |
| Tnfsf13b | 1 | 0.17 | 2.40 | 0.27 | 1.07 | 0.04 | 10.11 | 1.54 | 4.61 | 0.25 | No | No | Yes | No | Yes |
| Vegfa | 1 | 0.31 | 0.50 | 0.07 | 0.84 | 0.19 | 0.67 | 0.09 | 1.04 | 0.36 | No | No | No | No | No |
| Xcl1 | 1 | 0.24 | 8.08 | 0.64 | 3.67 | 0.58 | 16.19 | 3.12 | 36.19 | 7.80 | No | No | No | Yes | Yes |
