## Supplementary material for "Non-Necroptotic Roles of MLKL in Diet-Induced Obesity, Liver Pathology, and Insulin Sensitivity: Insights from a High Fat, High Fructose, High Cholesterol Diet Mouse Model": Table 3

| **Product** | **Control Diet (g)** | **Control Diet (kcal %)** | **HFHFrHC Diet (g)** | **HFHFrHC Diet (kcal %)** |
| --- | --- | --- | --- | --- |
| **Protein** | 19 | 20 | 22 | 20 |
| **Carbohydrate** | 67 | 70 | 45 | 40 |
| **Fat** | 4 | 10 | 20 | 40 |
| **Total** |  | 100 |  | 100 |
| **kcal/gram** |  | 3.8 |  | 4.5 |
| **Composition** | **Control Diet (g)** | **Control Diet (kcal)** | **HFHFrHC Diet (g)** | **HFHFrHC Diet (kcal %)** |
| **Protein** |  |  |  |  |
| Casein | 200 | 800 | 200 | 800 |
| L-Cystine | 3 | 12 | 3 | 12 |
| **Carbohydrate** |  |  |  |  |
| Corn Starch | 350 | 1400 | 0 | 0 |
| Maltodextrin 10 | 85 | 340 | 100 | 400 |
| Fructose | 0 | 0 | 200 | 800 |
| Dextrose | 169 | 676 | 0 | 0 |
| Sucrose | 96 | 384 | 96 | 384 |
| Cellulose | 50 | 0 | 50 | 0 |
| **Fat** |  |  |  |  |
| Soybean Oil | 25 | 225 | 25 | 225 |
| Primex Shortening | 0 | 0 | 0 | 0 |
| Palm Oil | 0 | 0 | 135 | 1215 |
| Lard | 20 | 180 | 20 | 180 |
| Mineral Mix S10026 | 10 | 0 | 10 | 0 |
| DiCalcium Phosphate | 13 | 0 | 13 | 0 |
| Calcium Carbonate | 5.5 | 0 | 5.5 | 0 |
| Potassium Citrate, 1H2O | 16.5 | 0 | 16.5 | 0 |
| Vitamin Mix V10001 | 10 | 40 | 10 | 40 |
| Choline Bitartrate | 2 | 0 | 2 | 0 |
| **Cholesterol** | 0 | 0 | 18 | 0 |

**Table S3**
